# Pan-cancer 3D genomic analysis revealed extremely long Polycomb loops as the biomarker for sensitivity to Polycomb inhibition

**DOI:** 10.1101/2025.10.30.685561

**Authors:** Sean Moran, Zhong Fan, Miriam Zanovello, Feng Fan, David Xue Qing Wang, Chao Lu, Jie Liu, Xiaotian Zhang

## Abstract

Polycomb targeted loci form long-range chromatin interactions independent of CTCF-cohesion and are demarcated by low DNA methylation(1–3). Polycomb targets form extremely long-range loops (long Polycomb loops) that can separate anchors up to 60 Mb. These loops predominantly occur in cells of self-renewal status, such as human hematopoietic stem cells (HSC) and mouse embryonic stem cells (ESC), but rarely in cell lines.

To identify long Polycomb loops in cancer, we initiated a pan-cancer survey of long Polycomb loops in a collection of 264 tumor samples (33 acute myeloid leukemias, 17 T cell lymphoblastic leukemia, 29 Breast cancer samples, 63 pediatric brain tumors, 70 prostate cancers, and 42 colon cancers). We found most cancers, including all prostate cancers and colon cancers, lack long Polycomb loops except pediatric brain tumors and certain AMLs.

In pediatric brain tumors, we found 30% of the ependymoma PFA subtype (which shows globally depleted H3K27me3) notably displayed strong long Polycomb loop interactions. AML displayed more diverse levels of long Polycomb loop interactions. Most AMLs lost both long Polycomb loops with Polycomb binding loss at the loop anchors. Whereas 10% of AMLs retain long Polycomb loops as strong as in HSCs. These AMLs recurrently carry mutations in *CEBPA* and *STAG2*, which are not associated with the Polycomb complex or DNA methylation machinery. We found that long Polycomb loop strong AML is sensitive to EZH2 inhibition, which induces cell differentiation. Conversely, PRC1 component dependency can be used to predict long Polycomb loop formation in B-cell lymphoblastic leukemia cell lines.

Our analysis suggests that the oncogenesis process antagonizes the long Polycomb loop maintenance in most cancers, yet certain cancers may still preserve strong long Polycomb loops from cell-of-origin. The maintenance of long Polycomb loops sensitizes cells to Polycomb inhibition, indicating that such loops could be an epigenomic biomarker for pharmacological or genetic Polycomb inhibition.

## Introduction

Eukaryotic genomes are folded into the 3D space of the nucleus through the formation of specific structures. During interphase, the genome forms distinct topologically associated domains (TADs), which represent the basic organizational unit of 3D chromatin. These megabase-sized structures are assembled by cohesin driven loop extrusion and the positioning of CTCF anchored barriers. Furthermore, chromosomal loops are formed between CTCF-occupied sites, whether they be TAD boundaries or intra-TAD CTCF anchors (4–7). Both TADs and loops are within 1 to 2 Mb in size. Their sizes are restricted by the density of CTCF/cohesin binding in the genome, which fluctuates around 20,000 to 40,000 in different cell types.

Besides TADs and loops, unidirectional cohesin loading can also lead to the formation of chromosomal stripes. Besides cohesin-CTCF, 3D genome structures mediated by other molecular mechanisms are relatively rare and occur in certain cell types at developmental stages or specific physiological and pathological processes (e.g., DNA replication and damage repair associated “fountain” and repair factory) (7–12). Among those 3D genome structures, 3D interactions between Polycomb target loci are common in the early development process from *Drosophila* to human (1,3,13–16)). In the mammalian genome, the interactions of Polycomb target loci appeared to display the following: (1). Long-range, where interactions formed between two loci separated by 20-60 Mb. (In *Drosophila*, interchromosomal interactions could also occur. This is due to the interacting loci being typical developmental genes which are sparse in the genome. (2). Independent of cohesin loop extrusion. (3). Early developmental features. (4). Dependent on both canonical PRC1 and PRC2 complexes. (5). The interacting loci are demarcated with Low DNA methylation between interacting loci across long genomic regions over 7.3kb (1,3,15,17). So far, disruption of PRC1 and PRC2 complex subunits by genetic and pharmacological means have led to the loss of long Polycomb loops (1,15–17). Meanwhile, disruption of cohesin complex members SCC1 enhances the long Polycomb interactions in mouse embryonic stem cells (mESCs)(18).

Details regarding the formation of Polycomb loops remain elusive due to the limited availability of high-resolution genome-wide Hi-C and imaging data sets. As such, the function of these long-range interactions in physiological and pathological conditions remains unclear.

The publication of several large cancer 3D genomic datasets has allowed us to systematically survey for long Polycomb loops in a pan-cancer setting (19–25). Across these disease models, the epigenomics of cancer often mimics early development, showing alterations of genome-wide Polycomb binding, global hypomethylation, and CpG island hypermethylation (26). Mutations that directly alter the Polycomb machinery, such as EZH2 gain-of-function mutations, Histone H3.3 K27M mutations, and overexpression of EZHIP protein, are frequently found in lymphoma and brain tumors. The long Polycomb loops dynamics across the different tumor types is unknown, given the high pan-cancer heterogeneity from tissue type and somatic mutations driving cancer.

In this study, we found that cancer cells displayed a reduction of long Polycomb loops, which was accompanied by a dramatic change in DNA methylation and polycomb loss. However, long Polycomb loops could still be preserved in tumor samples with recurrent somatic non-Polycomb complex protein mutations with sufficient Polycomb binding strength. We further identified tumors with long Polycomb loops that were addicted to the Polycomb machinery, with rare sensitivity to EZH2 inhibitors in their category of cancer.

Overall, we show that long Polycomb loops may be an epigenomic marker for cancers sensitive to polycomb disruption through small molecule inhibitors or genetic dependency of canonical PRC1 complex proteins.

## Result

### 1. Pan-Cancer Survey of long Polycomb loop

To study the long Polycomb loop (LPL) dynamics in cancer, we collected 264 publicly available Hi-C datasets from a spectrum of primary cancer samples and their corresponding normal tissues, including prostate cancer (n=79)(24), colon cancer of different stages (n=52)(20,21), breast cancer of different stages (n=29)(27), different subtypes of pediatric neurological cancer (n=63)(25), T cell lymphoblastic leukemia (n=17), and acute myeloid leukemia (n=33)(25). We applied aggregate peak analysis (APA) to tile up the interaction between all LPL anchors and called these LPLs (Figure 1A). DNA methylation canyon regions are stable across tissues, and those larger than 7.3 kb (referred to as Grand Canyons) reliably mark the strongest H3K27me3 domains in the genome. Because these domains form long-range 3D genomic interactions, we used Grand Canyon regions as genomic anchors for the LPL APA analysis (Figure 1A)(1). We used the conserved chromosomal loop regions defined before to perform quality control of pan-cancer samples first. Then, we used the LPL APA score to quantify the LPL interaction in the pan-cancer samples (Figure 1A, B).

**Figure 1.**
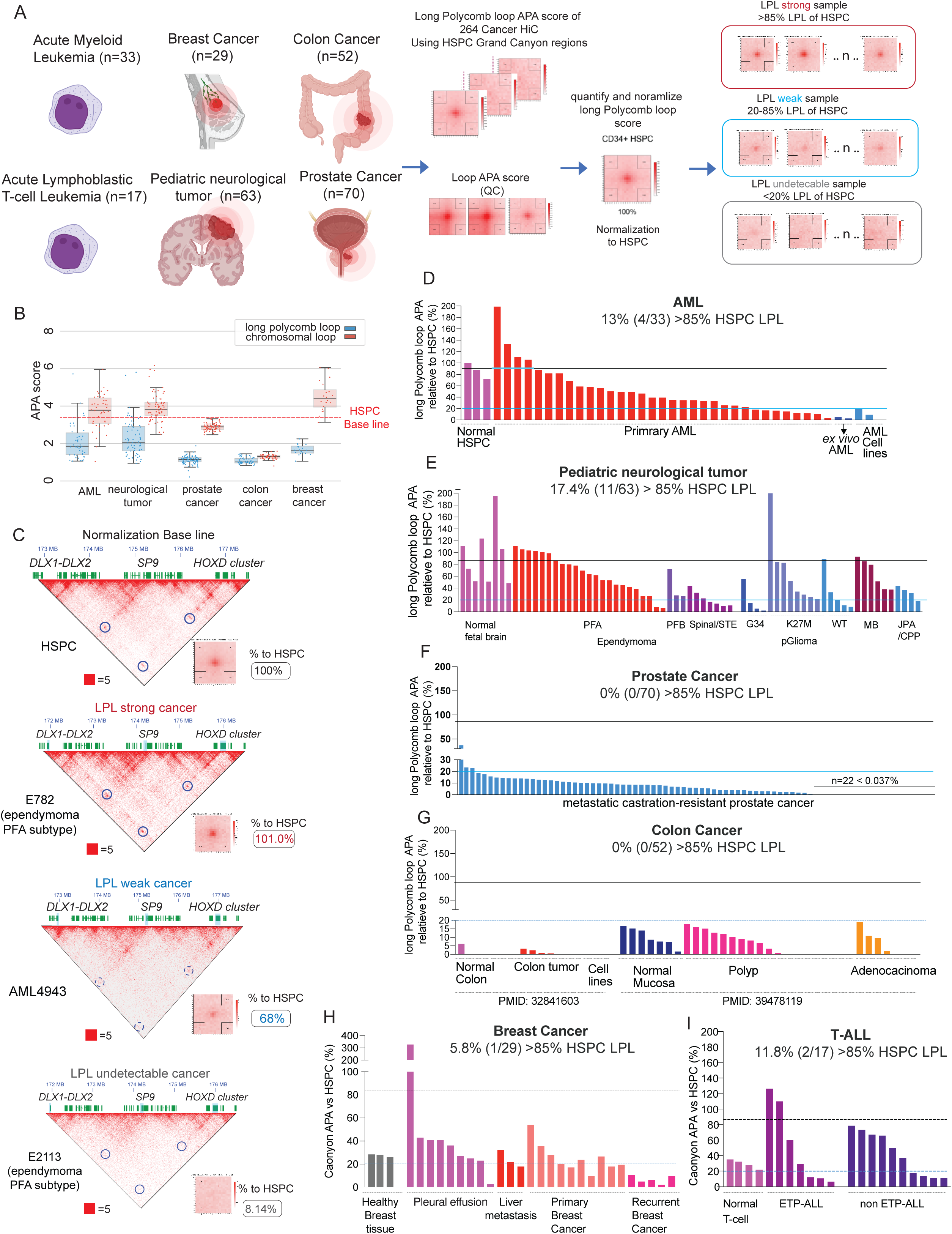
Pan-Cancer Survey of Long Polycomb Loop. **(A)** The strategy of long Polycomb loop pan-Cancer survey in over 200 HiC datasets. The LPL’s APA scores were used against the APA of a set of control loops for the quality control of the cancer Hi-C samples. The long Polycomb APA scores of cancer samples were normalized to the long Polycomb APA of the cord blood HSPCs. **(B)** The distribution of LPL APA score and loop APA score in the pan-cancer samples. A box plot of the 25th percentile, median, and 75th percentile of the APA score is shown. **(C).** The examples of LPL strong, weak, and undetectable cancer samples. The *HOXD* cluster – *SP9* – *DLX1/2* interactions from cancer samples containing different levels of LPL are shown for the visual inspection of LPL interaction strength. The accompanying long Polycomb loop APA plots are shown together with the HSPC normalized APA score level. (**D-I**). The waterfall plot of HSPC normalized LPL APA score distribution in AML (**D**), pediatric neurological cancer (**E**), prostate cancer (**F**), colon cancer (**G**), breast cancer (**H**) and T-ALL (**I**). LPL APA score of normal cells, tissue of the corresponding cancer was also included to compare with the cancer samples. The 85% HSPC LPL APA score (Black line) was added as a gauge for high LPL cancer samples. The 20% HSPC LPL APA score (Blue line) was added as the gauge for the detectable LPL interaction level. PFA: Posterior Fossa Group A, PFB: Posterior Fossa Group B, STE: supratentorial, MB: Medulloblastoma, JPA: Juvenile Pilocytic Astrocytoma, CPP: Choroid Plexus Papilloma.

We used HSPCs from cord blood as benchmark samples, as these contained strong LPL (Figure 1A, B). We observed that only a fraction of the samples contain LPL at a level stronger than HSPCs (Figure 1B). To categorize the pan-cancer tumor samples. We set 85% LPL APA in HSPCs as the threshold for “high LPL” (Figure 1A,C). In parallel, we have set the 20% LPL APA as the detection limit of LPL, as we were unable to visually detect the LPL signal in those samples (Figure 1A, C). Most of the samples are between 20 to 85% LPL, containing a visible LPL signal with a reduced level. In the examination of different types of cancers, we found that 13% (4 out of 33) of AML and 17.4% (11 out of 63) of pediatric neurological cancer samples contain high levels of LPL (Figure 1D, E, Supplemental Figure 1A-B). In contrast, in solid tumors, neither prostate nor colon cancer showed strong LPLs in solid tumors (Figure 1F, G, Supplemental Figure 1C-D). Most prostate and colon cancer samples lacked detectable levels of LPL (20% HSPC LPL APA score). This absence of LPLs was also observed across all types of breast cancers (Figure 1H, Supplemental Figure 1E). Interestingly, one single sample isolated from patients’ pleural effusion showed a high level of LPL (Figure1H). In pediatric neurological cancer samples, the posterior fossa type A (PF-EPN-A, PFA) ependymoma was the tumor subtype with strong LPL APA scores. 7 of 23 or 30.4%, of PFA, ependymoma showed a strong LPL APA score, the subtype with the greatest enrichment of high LPL tumors (Figure 1D). In contrast, the posterior fossa type B (PF-EPN-B, PFB), spinal and STE subtypes of ependymoma showed a low level of LPL. In other neurological tumors, H3K27M-mutated pediatric glioma and two medulloblastomas also showed strong LPL (Figure 1D). PFA ependymomas lack the genomic abnormalities, While PFA ependymomas have low global levels of H3K27me3, due to high EZHIP expression, they do possess strong localized H3K27me3 signals at canonical Polycomb targets. These strong Polycomb sites serve as anchors for LPLs. Additionally, we also observed that 10% of T-ALL samples displayed a high LPL signal, predominantly in the early T cell progenitor (ETP) subtype(Figure 1I). Since ETP-T-ALL’s cells of origin are very immature T-cell progenitors, the high LPL signal could be carried through the oncogenesis process. Overall, most of the cancer samples showed low or undetectable levels of LPL signal. Conversely, 10-30% of AML and pediatric neurological tumors displayed strong LPL signals, dependent on the disease subtype. There are two plausible explanations for the overall low level of LPL signal in pan-cancer samples. The first is that in cancer, the epigenome has been altered significantly at the LPL anchors – DNA methylation Grand Canyon regions – with DNA hypermethylation and Polycomb binding loss (26,28,29). Second, it is possible that the cell-of-origin from which the disease is derived already lacked LPLs. The strong LPL signals are observed in naive HSPCs and normal pediatric brain tissues, which are propagated to AML and pediatric neurological tumors (Figure 1D,E). As the tumor develops, the original strong LPL will likely be dampened by epigenomic reprogramming, leading to the loss of LPL in most fully developed tumors after the clonal expansion of the tumor cell of origin.

Interestingly, almost all the cell lines profiled show undetectable LPL (Figure.1 D,G), suggesting that the loss of LPL is associated with transformation and excessive cell proliferation in these cancer cell lines.

### 2. Recurrent transcriptomic and genomic signature of tumors with strong long Polycomb loop

Next, we wondered what the biomarker for the strong LPL signal in cancer could be. Both neurological brain tumors and AML have a relatively large sample size and comprehensive multiomics data. As PFA ependymomas lack somatic mutations or structural variations, we first performed transcriptomic analysis of the PFA ependymoma samples to find the gene signatures of LPL signal strength in PFA ependymoma. We analyzed the LPL APA correlated differentially expressed genes (DEGs) by ranking the PFA ependymoma samples’ LPL APA score. We found that the DEGs are clearly segregated into two distinct groups, as genes highly expressed or genes lowly expressed in LPL high samples (Figure 2A). We further analyzed gene ontology and found the lowly expressed genes in LPL low PFA ependymoma samples are enriched for neuronal differentiation associated protocadherin genes (Figure 2A), whereas the LPL high PFA ependymoma displayed the low differentiation phenotype and potential block of differentiation (Figure 2B). Next, we correlated DEGs with the sample’s LPL APA score. We observed that the homeotic transcription factor *MEIS1* expression positively correlates with the LPL APA score (Figure 2C). In contrast, Tuft cell-defining homeotic transcriptional factor - *POU2F*3’s expression anti-correlated with LPL APA score (Figure 2D). These data further show that a transcriptional program indeed drives the separation of LPL high and LPL low PFA ependymomas. As PFA ependymomas are known for the high expression of EZHIP and global loss but local enrichment of H3K27me3 at Polycomb target loci, we further correlated the PRC1 and PRC2 complex subunits’ gene expression with LPL APA score. We found that most of the PRC1/2 complex subunits are not correlated with the LPL APA score (Supplemental Figure 2 A-H). To our surprise, we found that the *EZHIP* gene, which drives PFA ependymoma development, is not correlated with the LPL APA score (Figure 2G). Other PRC1 complex subunits (*CBX2* and *PHC2*) were reported to form polymerization or condensate, but their expression is also not correlated with LPL APA score (Figure 2G). The expression level of *EZH1* and *PHF19* of the PRC2 complex significantly correlated with the LPL APA score (Figure 2E, F). But the fold change is milder from the LPL high to LPL low PFA ependymoma. Overall, the differentiation-associated transcriptional program is highly correlated with the LPL APA score in PFA ependymoma, suggesting the cell differentiation status could be another driving force for LPL interaction strength.

**Figure 2.**
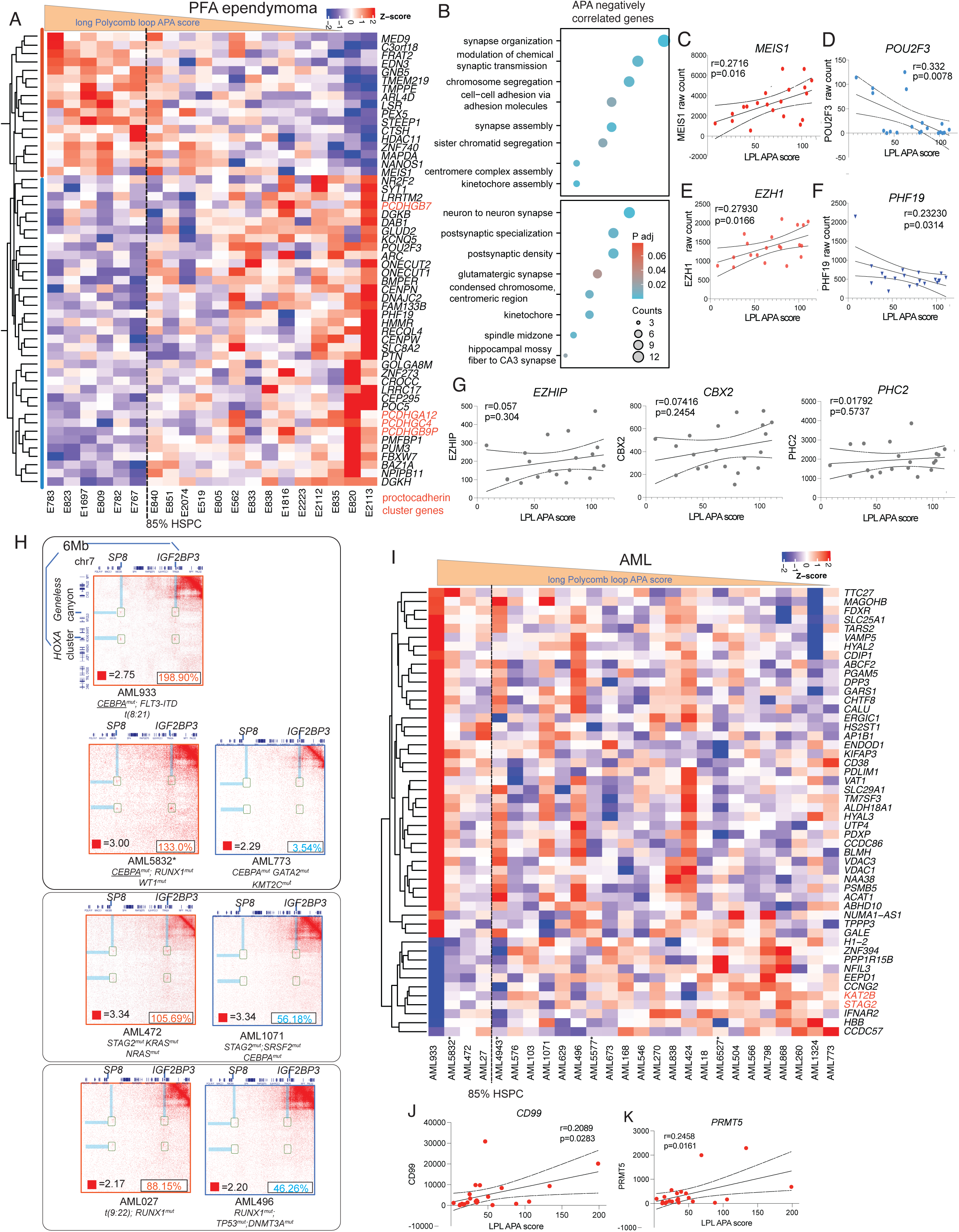
Transcriptomic and genomic signature of cancers with strong long Polycomb loop signal. **(A).** The top 50 differentially expressed genes associated with the LPL score in PFA ependymoma. The protocadherin genes are labelled. **(B).** The GO term analysis of genes whose expression level negatively correlated with the LPL APA score. (**C-D**). The Pearson correlation between expression of transcriptional factors – *MEIS1* (**C**), *POU2F3* (**D**), and LPL APA score in PFA ependymoma. The coefficient and p-value of the Pearson correlation are shown. (**E-F**). The Pearson correlation between expression of PRC2 subunit – *EZH1* (**E**), *PHF19* (**F**) and LPL APA score in PFA ependymoma. Pearson correlation analysis was performed. The coefficient and p-value of the Pearson correlation are shown. **(G).** The Pearson correlation between expression of *EZHIP, CBX2,* and *PHC2* correlated with LPL APA score in PFA ependymoma. The coefficient and p-value of the Pearson correlation were shown. **(H).** The 6Mb span of long Polycomb loops at the *HOXA* cluster to *SP8* from LPL high AML samples (left panel red box) and LPL low AML samples (right panel blue box). The genetic abnormalities of AML were annotated below the AML samples. **(I).** The top 50 differentially expressed genes associated with the LPL score in AML. *STAG2* and *KAT2B* are labelled. The AML samples generated by this study are marked by an asterisk. (**J-K**). The Pearson correlation between the expression of CD*99* (**J**), *PRMT5* (**K**), and LPL APA score in AML. The two genes show a positive correlation with the LPL APA score in AML. The coefficient and p-value of the Pearson correlation were shown.

Next, we performed the biomarker analysis for high LPL in AML. AMLs generally carry more than 3 somatic mutations per patient, and we first checked the somatic mutations LPL high AML carries, and we observed that recurring somatic mutations at *CEBPA* could be found in two LPL high AML from the combined two datasets (AML936 and AML5832) (Figure 2H). Besides mutations in *CEBPA*, mutations at *STAG2, and RUNX1* have also been observed in AML with a high level of LPL (Figure 2H). Furthermore, we analyzed for LPL APA score correlated DEGs in AML samples and found the gene expression clustering not as clear as the PFA ependymoma (Figure 2I). We could still observe that most of the DEGs in the LPL high and low groups are more enriched in the LPL high AML samples (Figure 2I). Also, two chromatin regulators – *KAT2B* and *STAG2* are lowly expressed in the LPL high AMLs. AML936 - the AML sample with the highest LPL signal, expressed the lowest level of *STAG2*. The data suggest that *STAG2* deficiency may contribute to the LPL maintenance during leukemogenesis. Previously, the deletion of cohesin subunit SCC1 was shown to enhance the LPL formation in mESCs (18), suggesting that a cohesin complex member might dampen the LPL formation during oncogenesis. The leukemia stem cell marker *CD99* is also positively correlated with LPL APA score (Figure. 2J).

### 3. DNA hypermethylation and Polycomb binding loss at long Polycomb loop anchors

Global DNA hypomethylation and CpG island (CGI) hypermethylation are hallmarks of cancer (26). Correspondingly, this leads to a loss of Polycomb at CGIs. At DNA methylation Grand Canyons, the anchor for long Polycomb loop, hypermethylation often occurs from the edge, resulting in a shrinking of the canyon region, loss of H3K27me3, and activation of local genes (28,29). To examine the epigenomic status at the anchors of long Polycomb loops, we comprehensively profiled whole genome bisulfite sequencing (WGBS), RNA-seq, H3K27me3, and H3K27ac in our in-house collection of acute myeloid leukemia (Figure 3A, Supplemental Table1). In these samples, AML5832 bears the strongest LPL (130% HSPC). AML4943 (*MLL-r*) bears a decreased level of LPL, but stronger than AML with DNA hypermethylation mutations - AML5577 (*IDH^R132H^*) and AML6527 (*TET^mut^*) (Figure 3B). Conversely, AML cell lines show a complete depletion of LPLs. 3D PCA clustering analysis showed LPL strong AML5832 clusters more closely to normal HSPCs, and other primary AMLs with medium/low LPLs clustered separately (Figure 3G). We compared the DNA methylation status at Grand Canyon (LPL anchor) and Canyon in primary AMLs to normal HSPCs. We found that DNA hypermethylation occurs in all the primary AML samples to the same level at both Grand Canyon and canyon regions (Figure.3C). While in AML cell lines, the strongest hypermethylation was observed at the Grand Canyon regions (Figure.3C), showing the gradual DNA hypermethylation that correlated with the level of LPL in oncogenesis transformation. Furthermore, the H3K27me3 level was compared across primary AMLs and HSPCs, and we found that in AML samples, the H3K27me3 level was decreased in all primary AMLs at Grand Canyon and canyons regions (Figure.3D). In LPL strong sample AML–AML5832, the H3K27me3 level was higher than two LPL low, DNA hypermethylated AML samples AML5577 and AML6527 (Figure 3D). This data suggests that there could be a threshold of low H3K27me3 level at LPL anchor regions to maintain the LPL. In the LPL medium sample AML4943, the H3K27me3 was also comparable to AML5832 (Figure 3D), further showing a general trend that a certain level of H3K27me3 is essential for LPL maintenance in cancer. We wondered whether, with the Polycomb mark loss in primary AML, the LPL anchor will be transcriptionally activated along with a gain in active H3K27ac mark. We found the Grand Canyon regions still lacked the H3K27ac mark with Polycomb mark loss (Figure 3E). We next examined an example locus at chromosome 2, across the *HOXD* cluster-*SP9*-*DLX1/2* region. In LPL strong AML5832, which shows a strong H3K27me3 distribution at the LPL anchor, also displays strong H3K27me3 spreading into surrounding genomic regions. In contrast, the DNA hypermethylated AMLs show a significantly lower H3K27me3 signal at both the LPL anchor loci and the surrounding genomic region (Figure 3F). Overall, the data shows epigenomic alterations in cancer – DNA hypermethylation and loss of H3K27me3 at LPL anchors (DNA grand canyons) are associated with loss of LPL during tumor development if the cell-of-origin contains LPL. Yet, in some cancers, the H3K27me3 level is decreased but remains sufficiently high to support the LPL formation.

**Figure 3.**
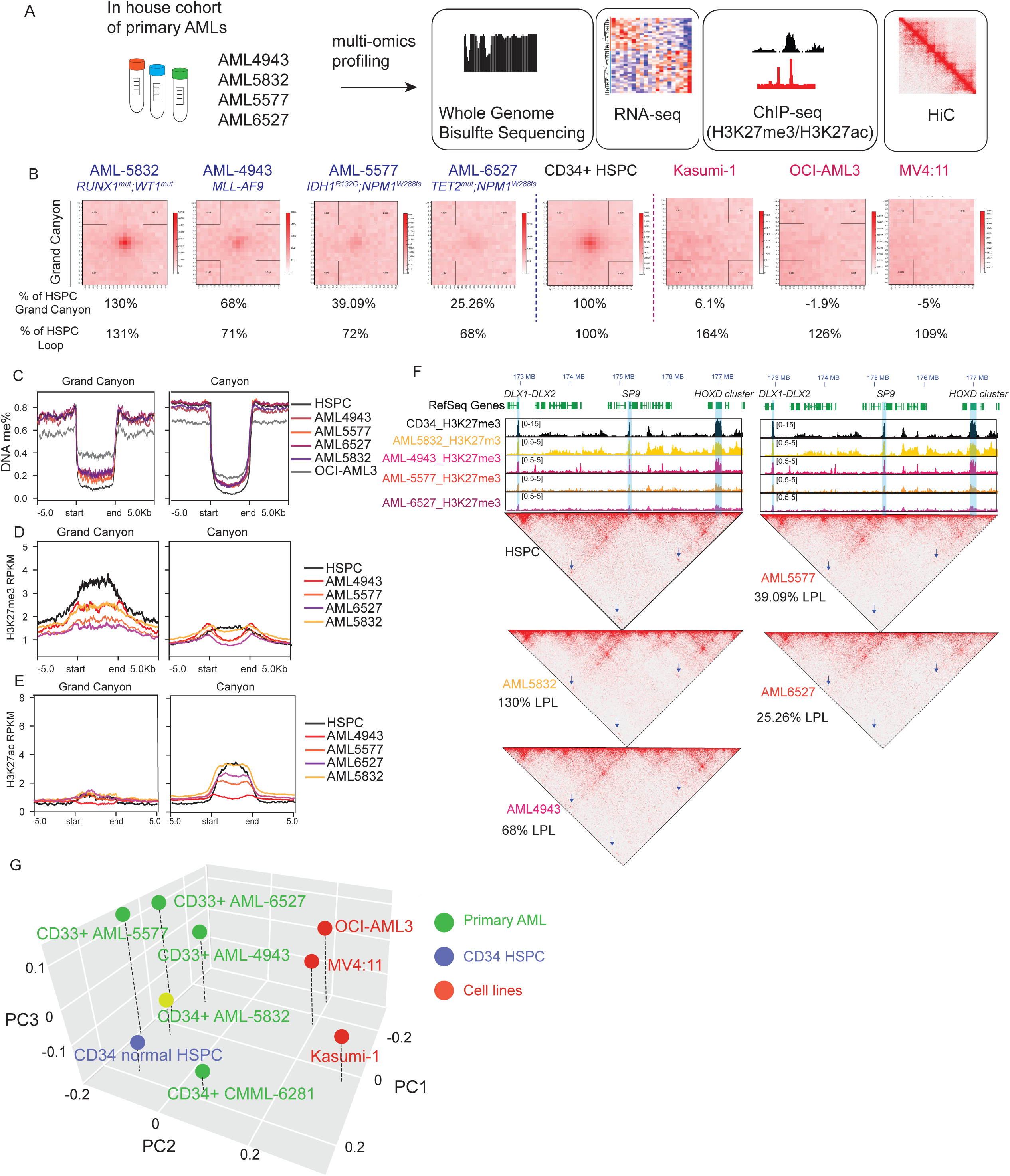
Long Polycomb loop anchors were hypermethylated and lost enrichment of H3K27me3 in cancer. **(A).** Scheme of multi-omics performed for AML samples used in this study **(B).** The LPL APA of the AML samples and cell lines used in this study. The LPL and loop APA scores of AML samples and cell lines are normalized to those of HSPC and are shown below the respective sample. **(C)** The DNA methylation metaplot of the Grand Canyon and Canyon regions in primary AML samples and the OCI-AML3 cell line. **(D).** The H3K27me3 metaplot of Grand Canyon and Canyon regions in primary AML samples. **(E).** The H3K27ac metaplot of Grand Canyon and Canyon regions in primary AML samples. **(F).** The H3K27me3 distribution at LPL anchored at *DLX1/2-SP9-HOXD* cluster regions. The HiC contact map is shown below. The arrow points to the LPLs. LPL anchors *DLX1/2, SP9,* and *HOXD* cluster were highlighted. **(G).** The three-dimensional clustering of primary AML samples and AML cell lines used in this study.

### 4. Long Polycomb loop is an epigenomic marker for Polycomb protein inhibitor sensitivity in cancer

We and others previously found EZH2 inhibition could disrupt the formation of LPL in normal HSPCs (1,15). We therefore wondered how cancers with strong LPL would respond to Polycomb complex targeting, such as EZH2 inhibition, given the potential LPL disruption. We performed EZH2 inhibition in LPL strong AML5832 cells. In these cells, we could observe AML-specific LPL in comparison with normal HSPCs (Figure 4A). Since AML5832 blast cells are CD34+, we isolated the CD34+ EZH2 enzymatic inhibitor – Tazemetostat (Figure. 4B). The AML5832 CD34+ cells showed dose-dependent sensitivity to Tazemetostat with a decrease in colony formation unit (CFU) in comparison with vehicle. Furthermore, we also observed morphological differentiation induced by the EZH2 inhibitor (Figure. 4C, D). H3K27me3 level also decreased globally after the Tazemetostat treatment (Figure. 4E). We then performed the RNA-seq in cells after the CFU formations and differential gene analysis after treatment. Cell cycle and DNA replication genes were strongly down-regulated, while immune response and phagosome associated genes were up-regulated (Figure 4F, G). The macrophage surface marker CD11b (*ITGAM*) and CD14 were strongly up-regulated, indicating cellular differentiation following Tazemetostat treatment (Figure 4G). Furthermore, we found a significant number of LPL anchored genes that were both up and down-regulated following treatment (Figure 4H). Canonical Polycomb target genes, such as *CDKN2B,* were upregulated (Supplemental Figure 3C,F). Repressed genes like *LHX9* were also activated at low levels (Supplemental Figure 3 A, D). Interestingly, we observed that LPL anchor located genes with basal level expression like *DLX1* and *HOXA6* were significantly down-regulated with H3K27me3 reduction (Figure 4H, Supplemental Figure 3 B,E). This down-regulation of expressed LPL anchored genes following EZH2 inhibitor treatment was previously observed at *HOXA* genes in normal HSPCs (1). The sum of this data suggests that high LPL could be an epigenomic marker for the sensitivity to Polycomb protein disruption. Such sensitivity to EZH2 enzymatic inhibition has rarely been seen in human primary AML, as well as the concomitant reduction in colony formation and cell differentiation enhancement To further validate the connection between Polycomb complex protein dependency and LPL interaction, we explored the DepMap database for the top cell lines with strong dependency on Polycomb complex 1/2 proteins. By using a gene effect cut-off of −1, we found that most Polycomb complex proteins have no significant effect on cell growth and survival. However, we found one B-cell acute lymphoblastic leukemia line, RCH-ACV, is highly sensitive to CRISPR knockout of PRC1 component PHC2 and CBX2 (Figure.4I-J). These two proteins were essential for the LPL formation in H3K27M gliomas (16). The Hi-C contact maps of RCH-ACV and its corresponding control set of cell lines RS4:11 and SEM were generated as all three cell lines contain the *NSD2* pE1099K mutation (22). We performed LPL APA analysis on all three samples and found the highest LPL signal in RCH-ACV cells (∼30% of HSPC signal) (Figure.4K) (22). In contrast, the RS4;11 and SEM cell lines displayed undetectable LPL scores (Figure.4K). In the examination of LPL, we found RCH-ACV showed robust LPL between canonical Grand Canyon regions such as *DLX1/2*-*SP9*; *HOXD* cluster-*SP9* and *HOXA* cluster-*IGF2BP3*; *Geneless Canyon* – *IGF2BP3* (Figure.4L,M). However, the global LPL APA score decreased due to loss of LPLs at regions such as the *HOXAD* cluster-*DLX1/2*. Interestingly, RS4;11 also showed long-range interaction at the *HOXA* cluster spanning ∼9 Mb (Figure.4M). However, such long interactions were not LPL in origin, but rather an enhancer-promoter interaction connecting the *HDAC9* region to the active segment of the *HOXA* cluster and the *SKAP2* gene (Supplemental Figure.3G). This data further indicates that the epigenomic status of the loop, rather than the size of the loop (distance between two anchors), is more functionally related to the PRC1/2 complex disruption sensitivity. Additionally, RCH-ACV also showed the strongest H3K27me3 enrichment across Grand Canyon regions (Figure. 4N).

**Figure 4.**
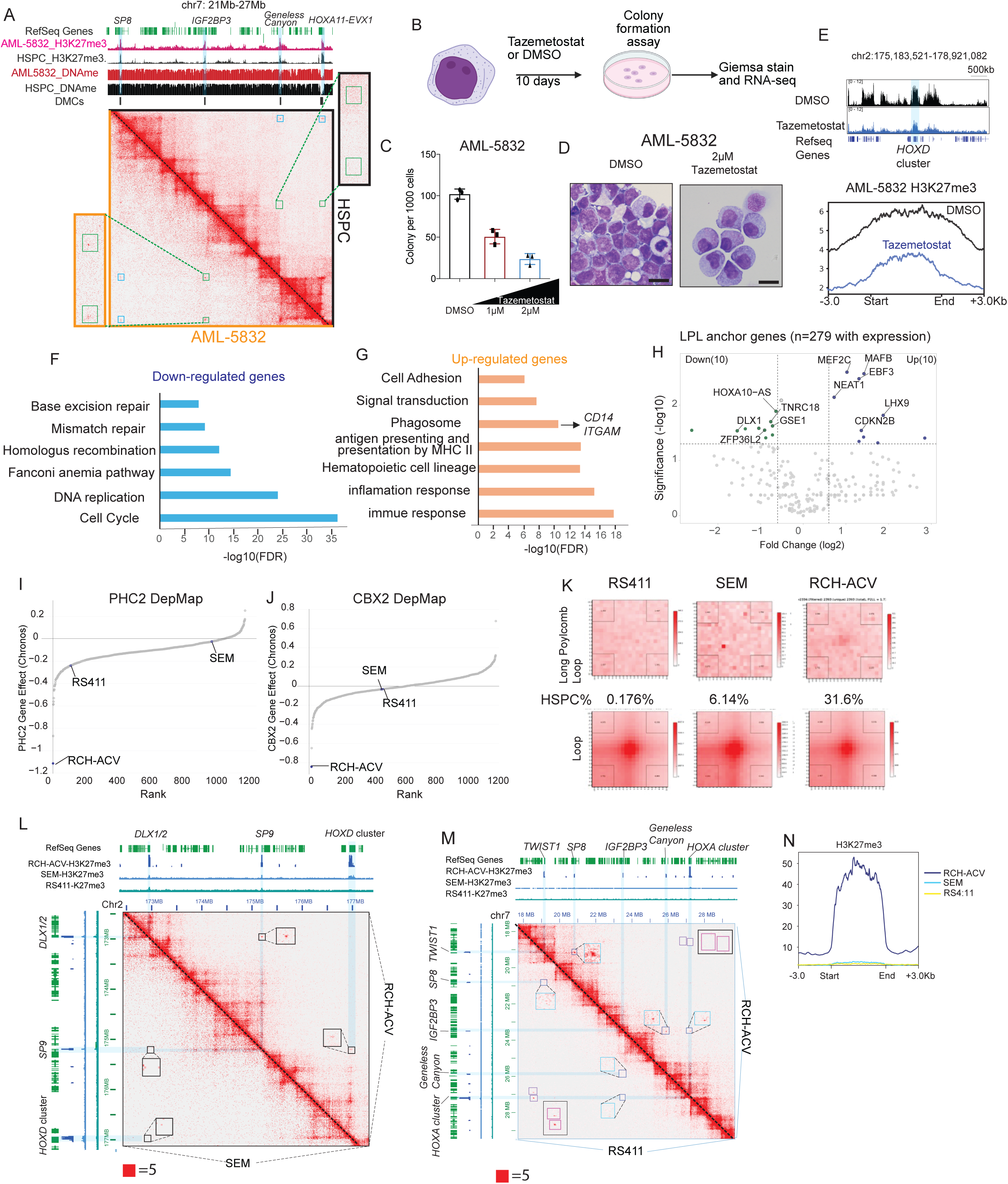
Correlation between sensitivity to PRC1/2 complex inhibition and long Polycomb loop strength in cancer. **(A).** The AML-5832 specific LPL interaction. The orange box zooms in on the AML-5832 specific interaction, which is lacking in HSPCs, zoomed in with blue boxes. DNA methylation and H3K27me3 of AML-5832 and HSPC, together with DNA methylation canyon (DMC), were shown. **(B).** EZH2 inhibitor Tazemetostat treatment scheme for AML5832 using colony formation assay. **(C).** The colony formation unit assay of AML5832 with an escalated dose of Tazemetostat. Mean + S.D was shown. **(D).** The Giemsa-wright staining of AML5832 cells treated with vehicle (DMSO) and 2μM Tazemetostat. Scale bar = 10μm. **(E).** The global reduction of H3K27me3 after the Tazemetostat treatment. **(F-G).** The gene ontology analysis of differentially expressed genes (F), Down-regulated genes **(G).** Up-regulated genes. **(H).** Volcano plot of differentially expressed LPL anchor genes. A fold change of 1.5 and a p-value of 0.05 was used as a cut-off. **(I-J).** DepMap dependency of PHC2 (I) and CBX2 (J). *NSD2 ^pE1099K^* mutated B cell lymphoblastic leukemia cell lines RCH-ACV, SEM and RS4:11 were labelled. **(K).** LPL and chromosomal loop APA analysis of *NSD2 ^pE1099K^*mutated cell lines RCH-ACV, SEM and RS4:11. **(L-M).** LPL in RCH-ACV cell lines at *DLX1/2-SP9-HOXD* cluster loci (**L**), and *TWIST1-SP8*; *IGF2BP3 - Geneless Canyon - HOXA* cluster loci (**M**). Boxed region shows the zoomed-in view of LPL at RCH-ACV and SEM HiC contact map in (**L**). Blue boxed region shows the LPL at RCH-ACV and RS4:11 contact map in (**M**). Red boxed region shows the long H3K27ac marked interaction at HOXA cluster in RS4:11 cell lines in (**M**). The LPL anchors are highlighted in blue. (**N**). H3K27me3 distribution metaplot over Grand Canyon regions in *NSD2 ^pE1099K^* mutated cell lines RCH-ACV, SEM and RS4:11.

### 5. Long Polycomb loop anchor genes are <u>derepressed</u> and act as novel cancer-specific domain organizers

Most cancers lack LPLs either because the cells-of-origin did not have LPL or the epigenomic alterations during oncogenesis erased the LPL over time. Little is known about the fate of LPL-repressed genes following these epigenomic changes in cancer cells. Previously, it was shown that genes located in these canyons became transcriptionally active due to hypermethylation (28)). We analyzed both our AML cohort and the TCGA AML cohort and found that the genes located within Grand Canyon LPL anchors were widely activated (Supplemental figure 4A, B). High HOX gene expression is a hallmark of many common subtypes of AMLs (*MLL-r* and mutant *NPM1*). We found a set of LPL anchor Grand Canyon genes – *IRX3, IRX5, NKX2-3,* and *FOXC1* in mutant *NPM1* AML and *HMX3* in MLL-r AML (Supplemental Figure 4B). These genes also correlated with *HOXA9* gene expression (Supplemental Figure 4B). We therefore hypothesized that the activated TFs may alter the 3D genomic architectures in cancer. We performed differential H3K27ac FitHiChIP analysis comparing AML and normal HSPC (Figure 5A). We categorized the H3K27ac loops into three *cis*-interacting groups: (1) *cis* interactions between H3K27ac-marked loci specific to AML; (2). *cis* interactions between H3K27ac-marked loci specific to normal HSPC; (3). *cis* interactions between conserved H3K27ac marked loci shared by normal HSPC and AML (Figure.5A). We found in AML (primary samples - AML4943 and cell lines – MV411 and OCI-AML3) most of *de novo* AML H3K27ac *cis* interactions occurs between AML specific H3K27ac marker loci (Group 1) (Figure.5B). In contrast, the HSPCs specific *cis* interactions are enriched for group 2 and group 3 cis interactions (Figure. 5B). In the further analysis of TF binding sites that enriched at anchors of group 1 *cis* interactions, we could observe the LPL anchor located homeotic TF binding sites are enriched at anchors of group 1 *cis* interactions in *HOX* ^hi^ cell lines - MV411 and OCI-AML3 (Figure. 5C-E). To elaborate this observation in detail, we found one AML-specific interaction – the *cis* interactions between non-coding RNA *TEX41* and key myeloid TF *ZEB2* could only be observed in the *HOX* ^hi^ primary AML and cell lines (Figure 5F,G). We could observe that the *TEX41* transcriptional start site (TSS) region obtains a leukemia-specific, H3K27ac-marked enhancer (Figure 5F). Importantly, the HOXA9 and ectopically activated LPL anchor located TF – IRX3 binding could be observed at the *TEX41* TSS (Figure 5G). The potential involvement of CTCF binding could also be excluded as the CTCF binding is consistent between AML and normal HSPCs, and the *TEX41* TSS region lacks the CTCF motif (Figure 5 F,G). IRX3 and IRX5 are important mesoderm-neuronal developmental TFs; ectopically expressed *IRX3* in AML was shown to work together with HOXA9 to block AML cell differentiation (30). Previously, an obesity associated enhancer – *FTO* intron enhancer was shown to regulate the *IRX3* expression(31). We also examined the cis interactions at *FTO-IRX3-IRX5* loci. We observed an interesting switch in 3D genomic architecture from repressive *IRX3-IRX5* long Polycomb loop formation to active *FTO-IRX3-IRX5* long-range enhancer-promoter interactions in primary AML blasts, cell lines, and lineage-ambiguous pediatric leukemia (Supplemental Figure 4C-E) (32).

**Figure 5.**
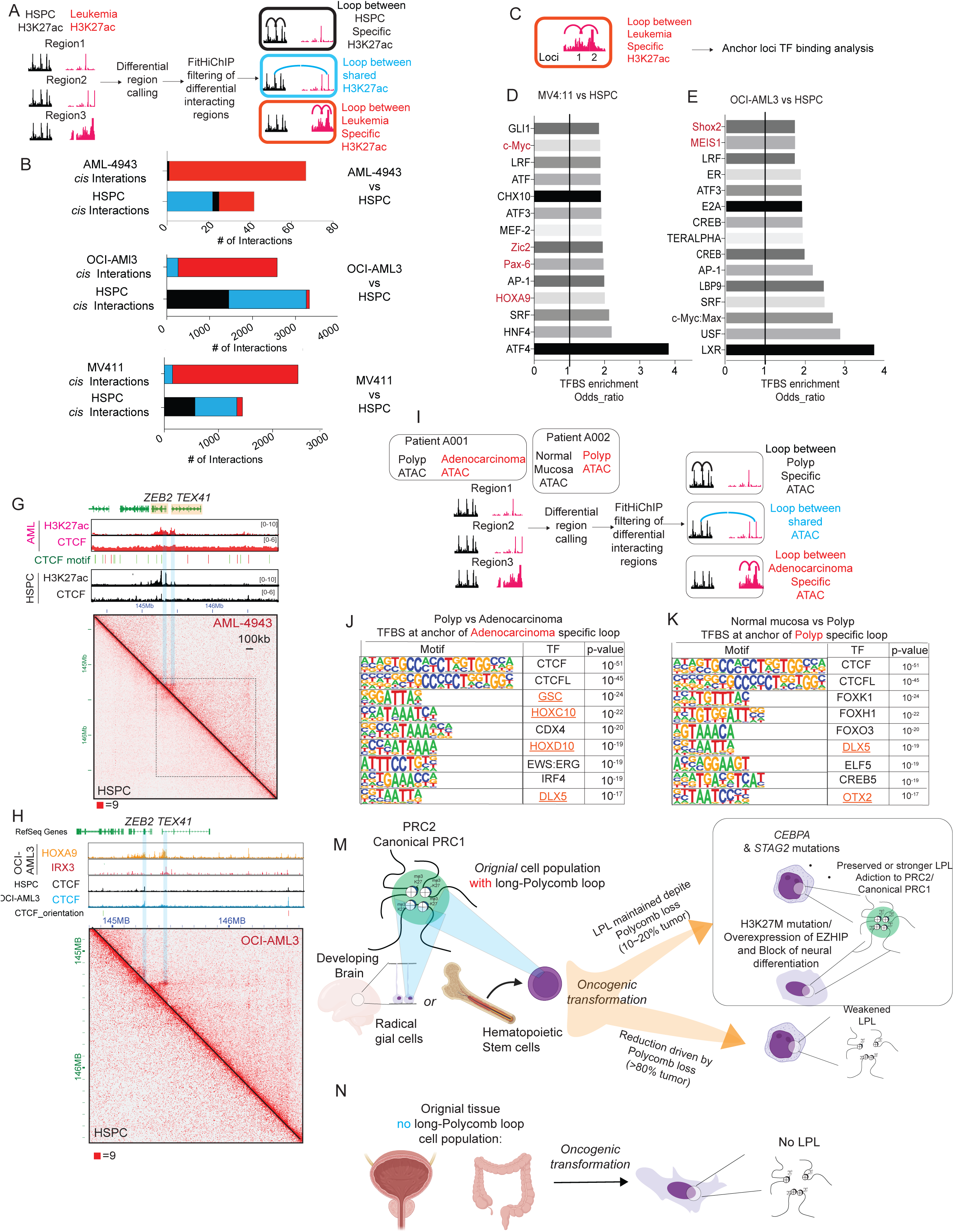
LPL anchor genes are activated and act as cancer-specific 3D genomic structures. **(A).** The scheme of differential *cis* interaction analysis using differential H3K27ac peak calling and FitHiChIP analysis in AML. **(B).** The composition of anchor regions for AML-specific *cis* interactions. Comparison of AML4943 vs HSPCs; OCI-AML3 vs HSPCs and MV4:11 vs HSPC was shown. **(C)**. TF binding analysis scheme for AML-specific *cis* interactions anchors from (**B**) (**D-E**). Motifs enriched in the leukemic-specific cis interactions in (**D**) MV411, (**E**) OCI-AML3. **(F-G).** The AML-specific cis interaction between ZEB2 and TEX41. AML4943 vs HSPC was shown in (**F**). The H3K27ac and CTCF were shown together in **F**. OCI-AML3 vs HSPC was shown in (**G)**. HOXA9, IRX3 binding in OCI-AML3 and CTCF in HSPCs and OCI-AML3 were shown in (**G**). **(H).** The scheme of differential *cis* interaction analysis using differential H3K27ac peak calling and FitHiChIP analysis in colon cancer. The paired polyp vs adenocarcinoma stages were compared using patient A001 data(20). The paired normal mucosa vs polyp stages were compared using patient A002 data (20). **(I-J).** Homer motif analysis of TF binding sites at A001 adenocarcinoma (**I**) and A002 polyp (**J**). The homeotic TFs located at LPL anchors (defined by DNA methylation Grand Canyon) are labelled orange. **(K-L).** The model of long Polycomb loop maintenance during carcinogenesis. (**K**). The cell-of-origin of PFA ependymoma in the developing brain and AML contains strong LPL. During oncogenesis, the loss of Polycomb will result in the weakened long LPL. In EPA ependymoma and H3K27M gliomas, the H3K27M and overexpression of EZHIP cluster the H3K27me3 and provide the prerequisite to maintain the LPL. The neuronal differentiation also erases LPL in PFA ependymoma. In AML, loss of *STAG2* and mutations of *STAG2* help to maintain the LPL. The somatic mutations in *CEBPA* may also contribute to LPL maintenance through self-renewal cell status maintenance. (**L**). The lack of long Polycomb loops in the cell of origin leads to low long Polycomb loop signals in colon and prostate cancer.

To further explore if the LPL anchor located TFs were activated and formed novel *cis* interactions in other cancers, we analyzed a colon cancer dataset containing paired normal-cancer HiC, ATAC-seq datasets from the same patient with the same strategy as in AML (Figure 5H). We performed a comparison between the pre-cancer polyp and normal colon (patient A001), as well as between the pre-cancer polyp and colorectal adenocarcinoma (patient A002). We found that the top TFs enriched in the differential *cis* interactions’ anchors specific to more cancerous stages are CTCF/CTCFL, suggesting the alterations could lead to topological domain alterations (Figure 5J, K). Interestingly, next to CTCF/CTCFL we found similar LPL anchor located TFs – DLX5 and OTX2 enriched in the polyp-specific *cis* interactions. As the polyp progresses to colorectal adenocarcinoma, more LPL anchor located TFs (*DLX5, GSC, HOXC10*, and *HOXD10*) are enriched in adenocarcinoma cis interactions (Figure 5I,J).

## Discussion

Overall, we found that in cancer, LPL interactions are, in general, undetectable and weak despite the self-renewal features of cancer. In AML and pediatric neurological tumors, 10 to 20% of patient samples show strong LPL interactions. In the pan-cancer setting, the cell-of-origin difference may explain the lack of LPL in solid tumor cancers like colon and prostate cancers (Figure 5K). In the cancer subtypes that cells-of-origin contain strong LPL interactions (HSCs and developing brain), LPL strength also reduces to a weak or undetectable level in most of the cancer samples. Therefore, the data suggest that the strong LPL signals are likely maintained but not created *de novo* during oncogenesis. The reduction in LPL could be explained by the epigenomic alteration in the LPL anchor – DNA methylation Grand Canyon during oncogenesis, particularly the reduction in Polycomb binding at the LPL anchors (Figure 5L). In 10-20% of cancer samples with strong LPL, certain somatic mutations on the epigenetic regulation pathway could create a permissible molecular mechanism for the maintenance of LPL. For example, the mutations in the PRC1/2 complex contribute to creating the permissible prerequisite for the formation. In PFA ependymoma, the overexpression of EZHIP could cluster the PRC1/2 protein and Polycomb mark-H3K27me3 at the LPL anchor by limiting the spreading of H3K27me3. These Polycomb enrichment mechanisms (overexpression of EZHIP or H3K27M mutation) could antagonize the Polycomb loss during oncogenesis and help to maintain the LPL. Yet, as we could also observe in the PFA ependymoma, the neuronal differentiation status can reduce the level of LPL, showing that Polycomb mutations are only essential but not sufficient for LPL maintenance (Figure 5K). Similar observations could be made in the AML cases, in which the *STAG2* mutation, or the low expression of *STAG2* – the non-Polycomb related mechanisms – could make LPL formation permissible. Again, such alterations are only essential but not sufficient for LPL maintenance in AML.

If the strong LPL is preserved in the cancer cells after the transformation, these LPL high cancer cells are sensitive to the targeting of PRC1/2 complexes through genetic or pharmacological methods. In AML and B-ALL cases, LPL high leukemias are sensitive to pharmacological or genetic inhibition of PRC1/2 complexes, particularly the inhibition of the canonical PRC1 complex subunit CBX2 and PHC2. These data suggest that a strong LPL signal could be an epigenomic marker for sensitivity to PRC1/2 inhibitors, such as EZH2 inhibitors, in pan-cancer settings. Currently, the clinical use of PRC1/2 inhibitors – primarily EZH2 inhibitors - is based solely on the somatic mutations that patients carry. These mutations include loss-of-function mutations in the SWI/SNF complex in epithelioid sarcomas and rhabdoid tumors, as well as gain-of-function mutations in EZH2 in lymphoma (33–36). Currently, no epigenome-based biomarker has been reported for the sensitivity of epigenetic targeting inhibitors. The sensitivity is predicted mostly based on genetic alterations, such as MATP synthetic lethality for PRMT5 inhibitors (37). LPL could be the first example of an epigenomic marker predicting the response to an epigenetic drug in cancer. Especially in AMLs and B-ALLs, the LPL high, EZH2 inhibitor sensitive samples do not carry the classic Polycomb or genetic biomarkers indicative of EZH2 inhibitor sensitivity. The data suggest the genomic markers may be insufficient to predict the PRC1/2 targeting. Therefore, the application of LPL as a biomarker for PRC1/2 targeting treatment could broaden the possibility for many tumors lacking a genomic-based biomarker for targeting therapy. The LPL dependence of these tumors may also provide the rationale to combine the EZH2 inhibitor with other commonly used cancer therapies for the enhancement of treatment.

Nevertheless, the mechanism sufficient to drive the formation of LPL remains unclear. The formation of LPL apparently requires a level of Polycomb mark over a certain threshold to enrich at the LPL anchors. However, the high level of Polycomb mark is just a prerequisite for LPL maintenance. Other mechanisms are likely to be present, enhancing the formation of LPL through the clustering of Polycomb marks, such as H3K27me3. Such LPL-enhancing mechanism likely involves the formation of the Polycomb body, which is now shown to be formed by the liquid-liquid phase separation (38,39). Through the formation of the Polycomb body, the interacting loci may also get enriched together in proximity, leading to the long-range interactions similar to those mediated by the NUP98 fusion protein (40). The post-translational modification (PTM) could further explain the mechanism through Polycomb body formation, as PTMs can drastically alter the condensate size and material properties, which can contribute to changes in Polycomb target locus interactions. Additionally, stem cell asymmetric division may be another process that contributes to the weakening of LPL maintenance. During stem cell asymmetric division in the fly testis, the differentiated daughter cells will lose the H3K27me3 during the asymmetric cell division that generates one stem cell and one differentiated cell (41). Such unequal inheritance of parental H3K27me3 histones and slow recovery of H3K27me3 in daughter cells will theoretically lead to the loss of H3K27me3 over time in differentiated populations. We also observed that the neuronal differentiation of PFA-EPN is strongly associated with the low or undetectable level of LPL (Figure. 2A). Considering the asymmetric division to symmetric division of radical glia progenitors is the initiating event of EPN(42), the symmetric division pattern to generate stem cell daughter cells may be a mechanism of long-term LPL maintenance besides the Polycomb sequestration.

## Materials and Methods

### Aggregate Peak Analysis for Canyons or Loops

The standard Aggregate Peak Analysis (APA) procedure was applied using Juicebox at 10 kb, except for colorectal cancer datasets, which were only available at 40 kb resolution. Juicer tools version 1.22.01 was used, and a master list of AML related methylation canyons was used to detect canyons, and a master list of AML associated chromatin loops was also used for assessing if a sample had viable Hi-C data.

All APAs were collected into comparative grids to assess the long Polycomb loop of specific datasets and of each sample.

### Thresholding for Long Polycomb loop calling

Long Polycomb loop relied on three general heuristics of increasing strictness to define whether a sample was (1) usable, (2) passed a minimum threshold of long Polycomb loop activity or not, and finally (3) if the activity of long Polycomb loop was considered to be high.

*(1) Criteria for Sample Usability:* To assess sample usability we perform an APA on the sample for the chromatin loops within “CD34pos” loops. High quality samples will have these loops fully visible in the aggregation, as both a clear spot in the loop APA and its APA score, typically above 2.0 but dependent on the provided dataset (i.e. samples were only 40 kb resolution was available will have another threshold value).
*(2) Minimum Criteria for Long Polycomb loop:* To be considered for long Polycomb loop the sample must first be (1) usable. Then, in the Canyon APA a spot should be visible, and the APA score should exceed 1.3.
*(3) Criteria for High Long Polycomb loop/Activity:* The threshold for high long Polycomb loop is satisfying (1) usability, (2) minimum criteria for long Polycomb loop, and finally the requirements outlined here. We take the ratio of the sample’s canyon-APA score to the control sample (1258-hic or GSM2861708). This ratio should be greater than or equal to 1.0. The raw APA-score should also be greater than 3.0, and visually inspecting the APA chart should show a clear red peak, not a reddish “shadow.”

### HiC matrix conversion and visualization

In cases that only mcool or ice-matrix formats were available, we used utility scripts for conversion. Dot-matrix files were converted into mcool using hicpro2higlass from HiC-Pro, and mcool or cool files were converted into hic-format using hicConvertFormat from HiCExplorer. Hi-C matrices for individual canyons were made using hic-straw, matplotlib, and seaborn. Track data was visualized using Deeptools’ PyBigWig library.

### Gene expression correlation analysis with long Polycomb score

#### Hi-C library generation for cell line and primary human AML

OCI-AML3, MV4-11 and Kasumi-1 cell lines Hi-C libraries were constructed with Arima Hi-C kit according to the manufacturer’s instruction. 1-2 million cells are used for the construction of Hi-C libraries. Each sample was sequenced to low depth to preform quality control check with Hi-C-Pro before sequencing to high depth.

Primary human AML samples were purchased from Stem Cell & Xenograft Core at University of Pennsylvania from de-identified frozen mononuclear cells from AML patients. Samples were obtained after IRB consent and provided to us as annotated, anonymous samples. Mononuclear cells were thawed by directly pipetting ice cold PBS+2% BSA to frozen cells in the freezing vials under room temperature. Thawed mononuclear cells were then gradient separated with Lymphoprep (StemCell Tech). Buffy coat was then collected to get the alive cells. Cells were stained for CD33 (PE, clone WM53, BioLegend), CD34 (APC, clone 581, BioLegend), CD45 (FITC, clone HI30, BioLegend), and CD38 (PeCy7, clone HB-7, BioLegend) to check for immunophenotyping by flow cytometry. Mononuclear cells were then fixed with 2% Formaldehyde for 10mins and quenched by 0.125M glycine for 5mins. *In situ* Hi-C is performed with fixed mononuclear cells following the previous protocol. Cells were digested by DpnII (NEB) for 2 hours, end filling with biotin-dATP (Jena Bioscience) and then ligated overnight. Ligated DNA was isolated by Pheno-chloroform extraction and ethanol precipitation with phase lock tube (Qiagen). Isolated DNA was then sonicated to 200-600bp with Covaris E220 and enriched for biotin containing ligated fragments using streptavidin magnetic beads (ThermoFisher). Enriched biotin containing ligated fragments were then made to library using Accel-NGS 2S kit (Swift Bioscience).

### Primary human AML *ex vivo* expansion and colony forming assay

AML5832 cells are thawed, and CD34+ cells were isolated using CD34+ selection beads (CD34 MicroBead Kit, human, Miltenyibiotec). CD34+ were recovered overnight after thawing in *ex vivo* expansion medium (SFEM II, 100ng/mL SCF, TPO, FLT3L, 35nM UM171, 1μM StemRegenin1 (SR1)). 10,000 AML5832 CD34+ cells were then plated in 6 well plate with 1mL of MethoCult™ H4034 Optimum (StemCell technology). Colonies were counted 10 days after plating and the colony pictures were taken at the same time. Drug treatment was performed using the EZH2 inhibitor (Tazemetostat, Selleckchem) at the indicated concentration.

### ChIP-seq

ChIP in cell lines, and primary samples are performed with previously described ChIPmentation protocol. Primary samples are centrifuged with Lymphoprep gradient (StemCell Tech) for the selection of alive cells. Alive cells are washed with PBS once and counted for the number and viability. One million viable cells were fixed with 1% formaldehyde and sonicated with Covaris E220. Sonicated chromatin were then incubated with antibody against H3K27me3(Cell Signaling, #9733), CTCF (Abcam, #ab70303), or H3K27ac (Diagenode, C15410174) overnight. Protein A/G beads (ThermoFisher) were added to the sonicated chromatin for 2 hours. After washes with low-salt, high-salt and LiCl wash buffer. Pull-downed DNA was de-crosslinked and digested with Proteinase K (Zymo Research) at 55 °C for 3 hours. Pull-downed DNA was extracted by DNA clean and concentration kit (Zymo Research). Libraries were made using Accel-NGS 2S kit (Swift Bioscience (currently IDT)).

### Whole genome bisulfite sequencing

Genomic DNA was isolated using QIAamp DNA Mini Kit (Qiagen). After isolation, around 1ug of genomic DNA was bisulfite converted using the EZ DNA Methylation-Lightning kit (Zymo). Whole genome bisulfite sequencing libraries were generated using Accel-NGS 2S kit (Swift Bioscience) with bisulfite-converted DNA. After sequencing, Fastq files are aligned to the corresponding genomic assembly using Bismark. CpG sites less than 5 times were filtered out.

## Data availability

The pan-Cancer HiC datasets are collected from the following: AML (GSE152135 and GSE302258); pediatric neurological tumor (GSE186599); prostate cancer (GSE249494); breast cancer (GSE261230 and GSE261230); colon cancer (GSE133928 and GSE207954); B-ALL cell line (GSE199754).

The in-house AML datasets have been deposited under the following GEO records: GSE302061 for RNA-seq, GSE302129 for ChIP-seq, GSE302062 for WGBS, and GSE302258 for HiC.

## Acknowledgement

This work was supported by NIH funding: R01HL170115, AACR MPM translational oncology award, and UTHealth Academic Excellence Award to Xiaotian Zhang. Jie Liu was supported by NIH funding (U01DK135017, R35HG011279) during the project execution. Chao Lu was supported by NIH funding R01DE03187, R01CA266978, R01DK132251.

## Notes

### Competing Interest Statement

The authors have declared no competing interest.

### Summary of Updates

We changed the figure curation and text to improve the manuscript.

