## Supplemental data for "Pan-cancer 3D genomic analysis revealed extremely long Polycomb loops as the biomarker for sensitivity to Polycomb inhibition"

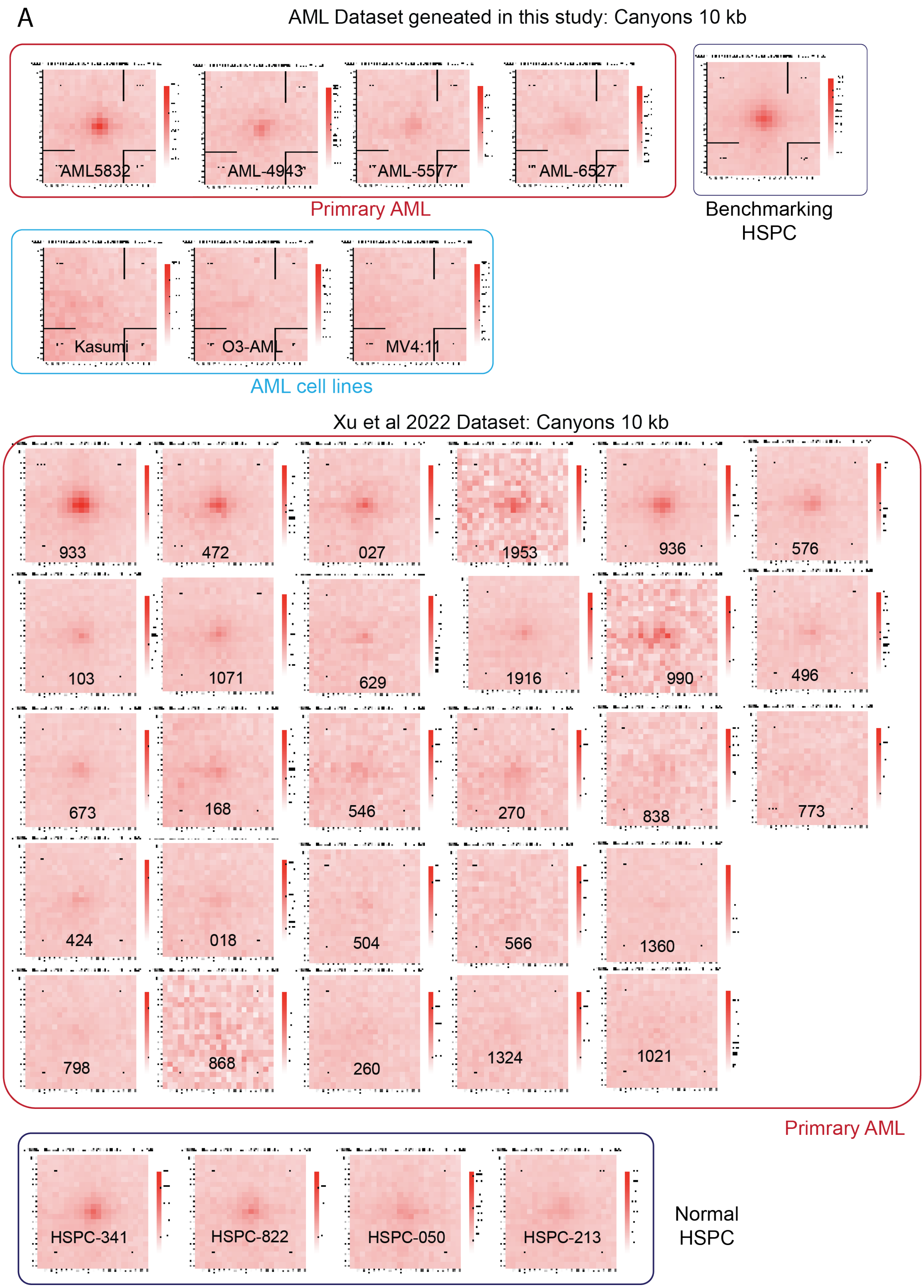
A
AML Dataset geneated in this study: Canyons 10 kb
AML5832
AML-4943
AML-5577
AML-6527
Primrary AML
Benchmarking HSPC
Kasumi
O3-AML
MV4:11
AML cell lines
Xu et al 2022 Dataset: Canyons 10 kb
933
472
027
1953
936
576
103
1071
629
1916
990
496
673
168
546
270
838
773
424
018
504
566
1360
798
868
260
1324
1021
Primrary AML
HSPC-341
HSPC-822
HSPC-050
HSPC-213
Normal HSPC

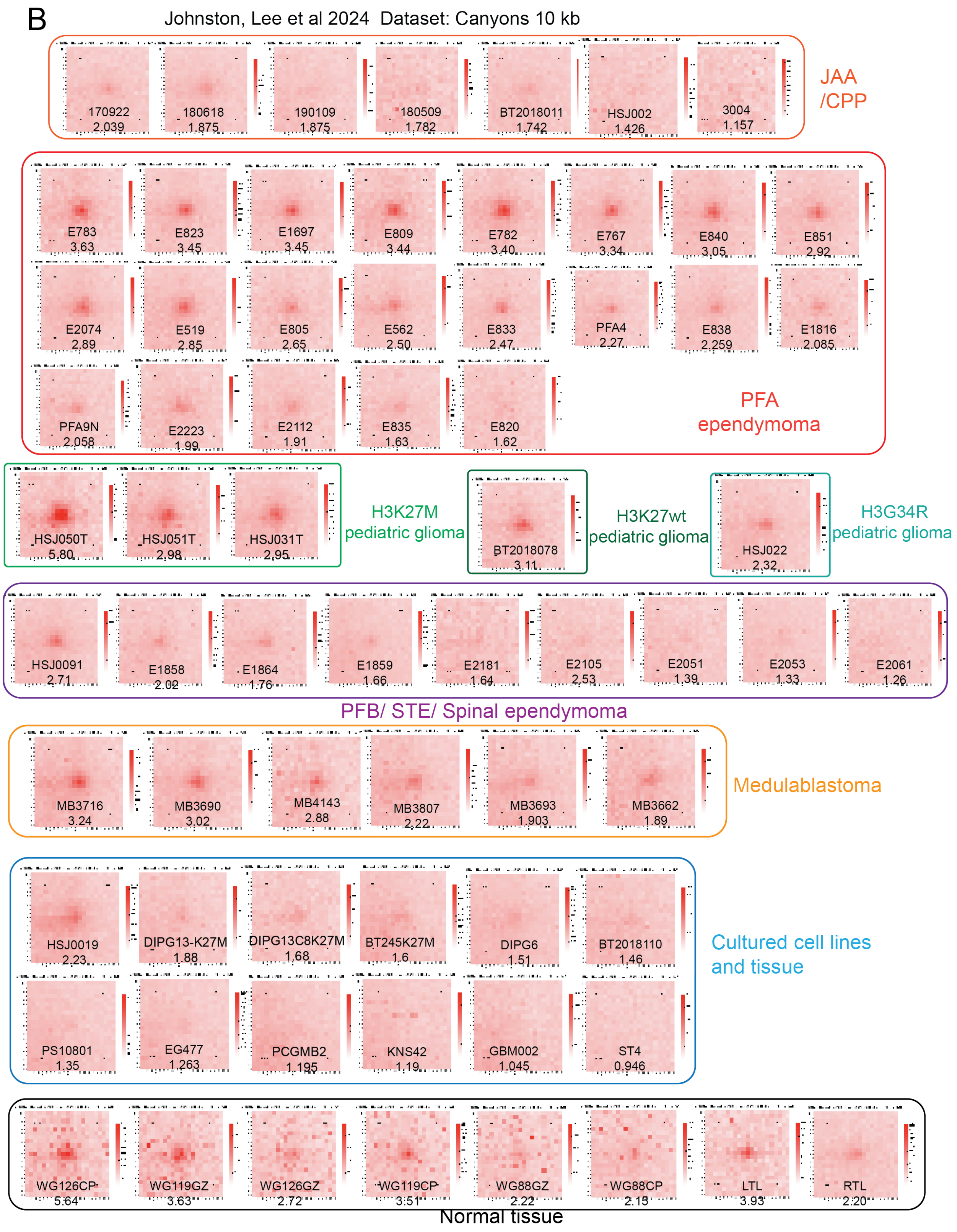
B
Johnston, Lee et al 2024 Dataset: Canyons 10 kb
170922 2.039
180618 1.875
190109 1.875
180509 1.782
BT2018011 1.742
HSJ002 1.426
3004 1.157
JAA /CPP
E783 3.63
E823 3.45
E1697 3.45
E809 3.44
E782 3.40
E767 3.34
E840 3.05
E851 2.92
E2074 2.89
E519 2.85
E805 2.65
E562 2.50
E833 2.47
PFA4 2.27
E838 2.259
E1816 2.085
PFA9N 2.058
E2223 1.99
E2112 1.91
E835 1.63
E820 1.62
PFA ependymoma
HSJ050T 5.80
HSJ051T 2.98
HSJ031T 2.95
H3K27M pediatric glioma
BT2018078 3.11
H3K27wt pediatric glioma
HSJ022 2.32
H3G34R pediatric glioma
HSJ0091 2.71
E1858 2.02
E1864 1.76
E1859 1.66
E2181 1.64
E2105 2.53
E2051 1.39
E2053 1.33
E2061 1.26
PFB/ STE/ Spinal ependymoma
MB3716 3.24
MB3690 3.02
MB4143 2.88
MB3807 2.22
MB3693 1.903
MB3662 1.89
Medulablastoma
HSJ0019 2.23
DIPG13-K27M 1.88
DIPG13C8K27M 1.68
BT245K27M 1.6
DIPG6 1.51
BT2018110 1.46
PS10801 1.35
EG477 1.263
PCGMB2 1.195
KNS42 1.19
GBM002 1.045
ST4 0.946
Cultured cell lines and tissue
WG126CP 5.64
WG119GZ 3.63
WG126GZ 2.72
WG119CP 3.51
WG88GZ 2.22
WG88CP 2.13
LTL 3.93
RTL 2.20
Normal tissue

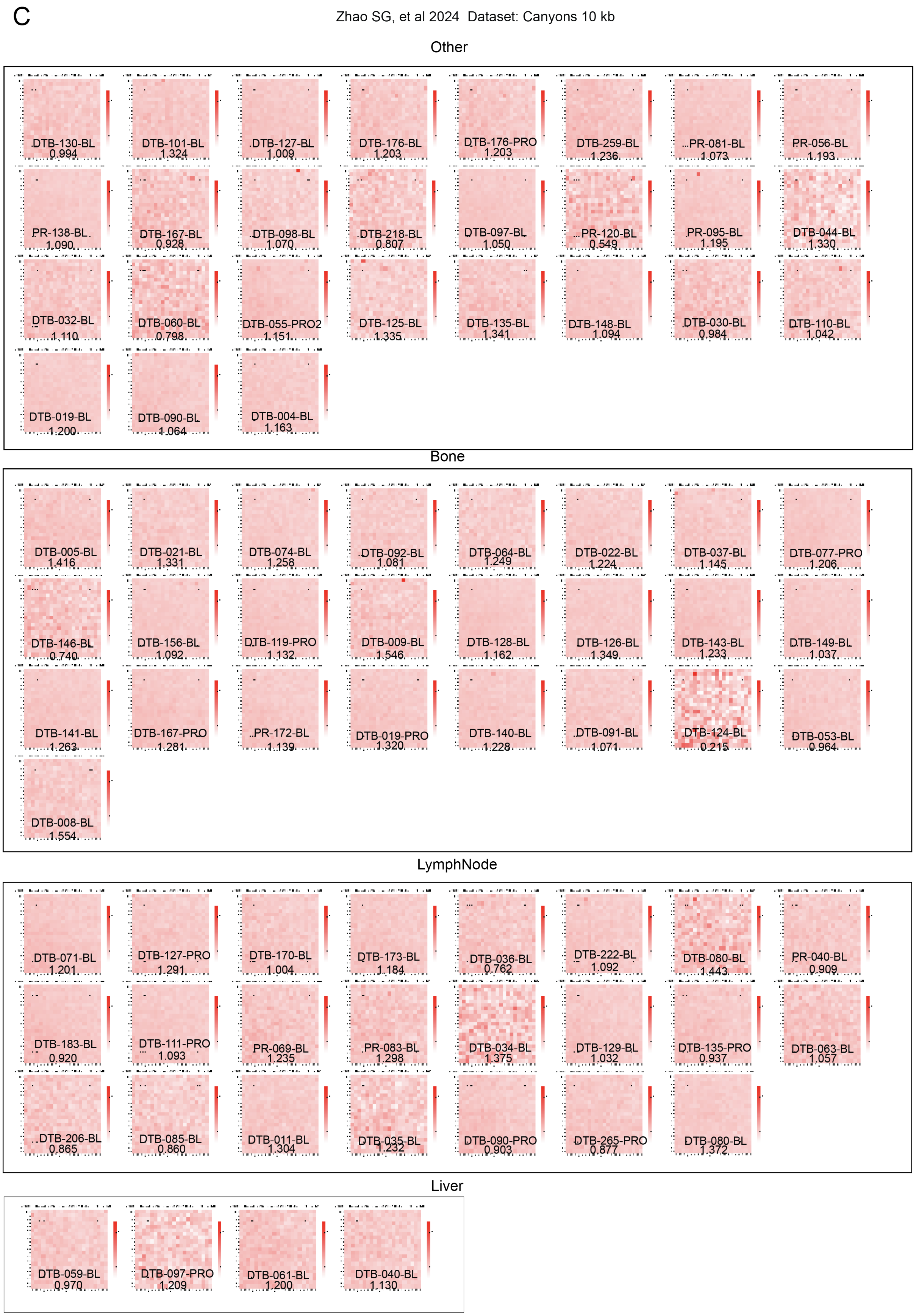
C
Zhao SG, et al 2024 Dataset: Canyons 10 kb
Other
DTB-130-BL 0.994
DTB-101-BL 1.324
DTB-127-BL 1.009
DTB-176-BL 1.203
DTB-176-PRO 1.203
DTB-259-BL 1.236
PR-081-BL 1.073
PR-056-BL 1.193
PR-138-BL 1.090
DTB-167-BL 0.928
DTB-098-BL 1.070
DTB-218-BL 0.807
DTB-097-BL 1.050
PR-120-BL 0.549
PR-095-BL 1.195
DTB-044-BL 1.330
DTB-032-BL 1.110
DTB-060-BL 0.798
DTB-055-PRO2 1.151
DTB-125-BL 1.335
DTB-135-BL 1.341
DTB-148-BL 1.094
DTB-030-BL 0.984
DTB-110-BL 1.042
DTB-019-BL 1.200
DTB-090-BL 1.064
DTB-004-BL 1.163
Bone
DTB-005-BL 1.416
DTB-021-BL 1.331
DTB-074-BL 1.258
DTB-092-BL 1.081
DTB-064-BL 1.249
DTB-022-BL 1.224
DTB-037-BL 1.145
DTB-077-PRO 1.206
DTB-146-BL 0.740
DTB-156-BL 1.092
DTB-119-PRO 1.132
DTB-009-BL 1.546
DTB-128-BL 1.162
DTB-126-BL 1.349
DTB-143-BL 1.233
DTB-149-BL 1.037
DTB-141-BL 1.263
DTB-167-PRO 1.281
PR-172-BL 1.139
DTB-019-PRO 1.320
DTB-140-BL 1.228
DTB-091-BL 1.071
DTB-124-BL 0.215
DTB-053-BL 0.964
DTB-008-BL 1.554
LymphNode
DTB-071-BL 1.201
DTB-127-PRO 1.291
DTB-170-BL 1.004
DTB-173-BL 1.184
DTB-036-BL 0.762
DTB-222-BL 1.092
DTB-080-BL 1.443
PR-040-BL 0.909
DTB-183-BL 0.920
DTB-111-PRO 1.093
PR-069-BL 1.235
PR-083-BL 1.298
DTB-034-BL 1.375
DTB-129-BL 1.032
DTB-135-PRO 0.937
DTB-063-BL 1.057
DTB-206-BL 0.865
DTB-085-BL 0.860
DTB-011-BL 1.304
DTB-035-BL 1.232
DTB-090-PRO 0.903
DTB-265-PRO 0.877
DTB-080-BL 1.372
Liver
DTB-059-BL 0.970
DTB-097-PRO 1.209
DTB-061-BL 1.200
DTB-040-BL 1.130

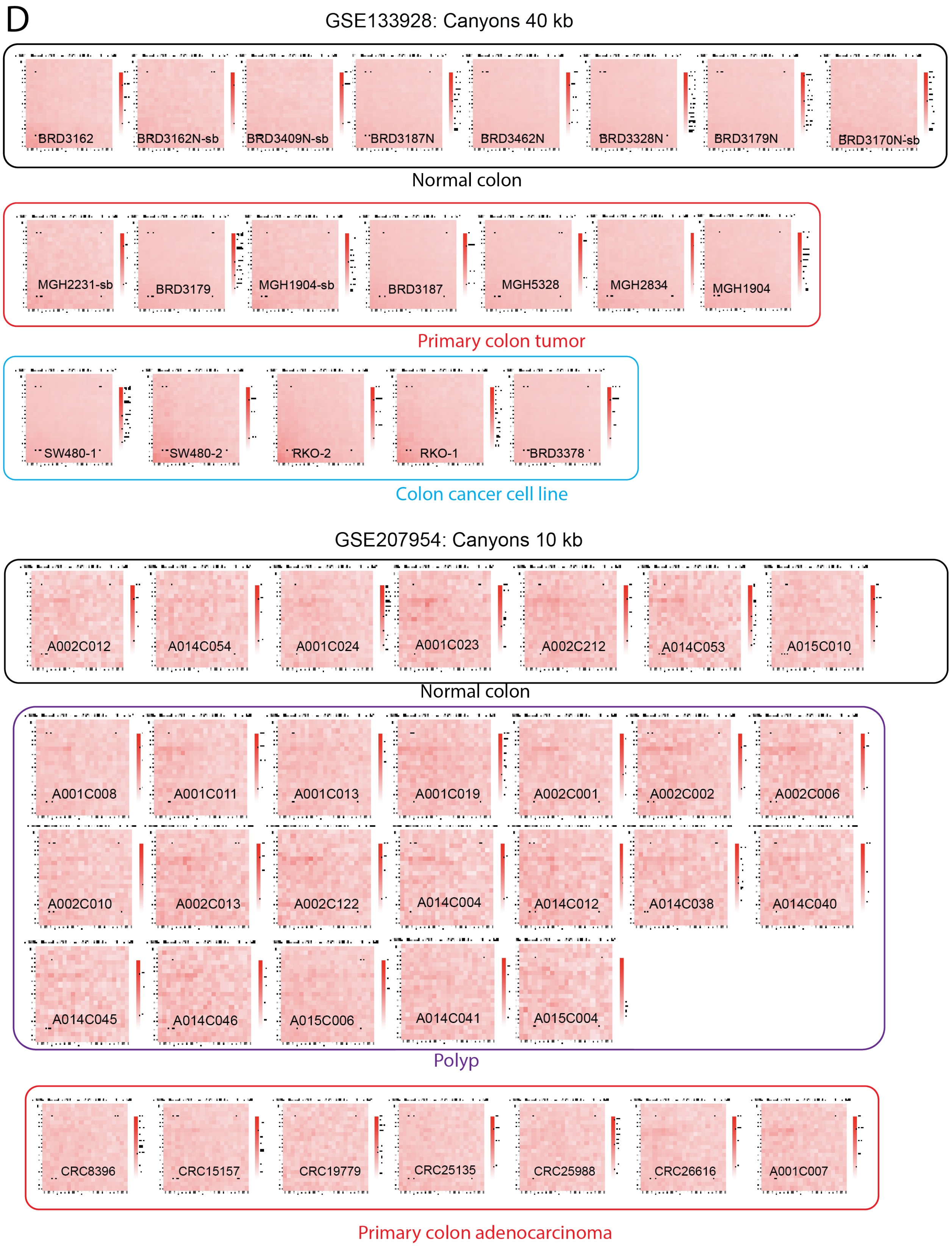
D
GSE133928: Canyons 40 kb
BRD3162
BRD3162N-sb
BRD3409N-sb
BRD3187N
BRD3462N
BRD3328N
BRD3179N
BRD3170N-sb
Normal colon
MGH2231-sb
BRD3179
MGH1904-sb
BRD3187
MGH5328
MGH2834
MGH1904
Primary colon tumor
SW480-1
SW480-2
RKO-2
RKO-1
BRD3378
Colon cancer cell line
GSE207954: Canyons 10 kb
A002C012
A014C054
A001C024
A001C023
A002C212
A014C053
A015C010
Normal colon
A001C008
A001C011
A001C013
A001C019
A002C001
A002C002
A002C006
A002C010
A002C013
A002C122
A014C004
A014C012
A014C038
A014C040
A014C045
A014C046
A015C006
A014C041
A015C004
Polyp
CRC8396
CRC15157
CRC19779
CRC25135
CRC25988
CRC26616
A001C007
Primary colon adenocarcinoma

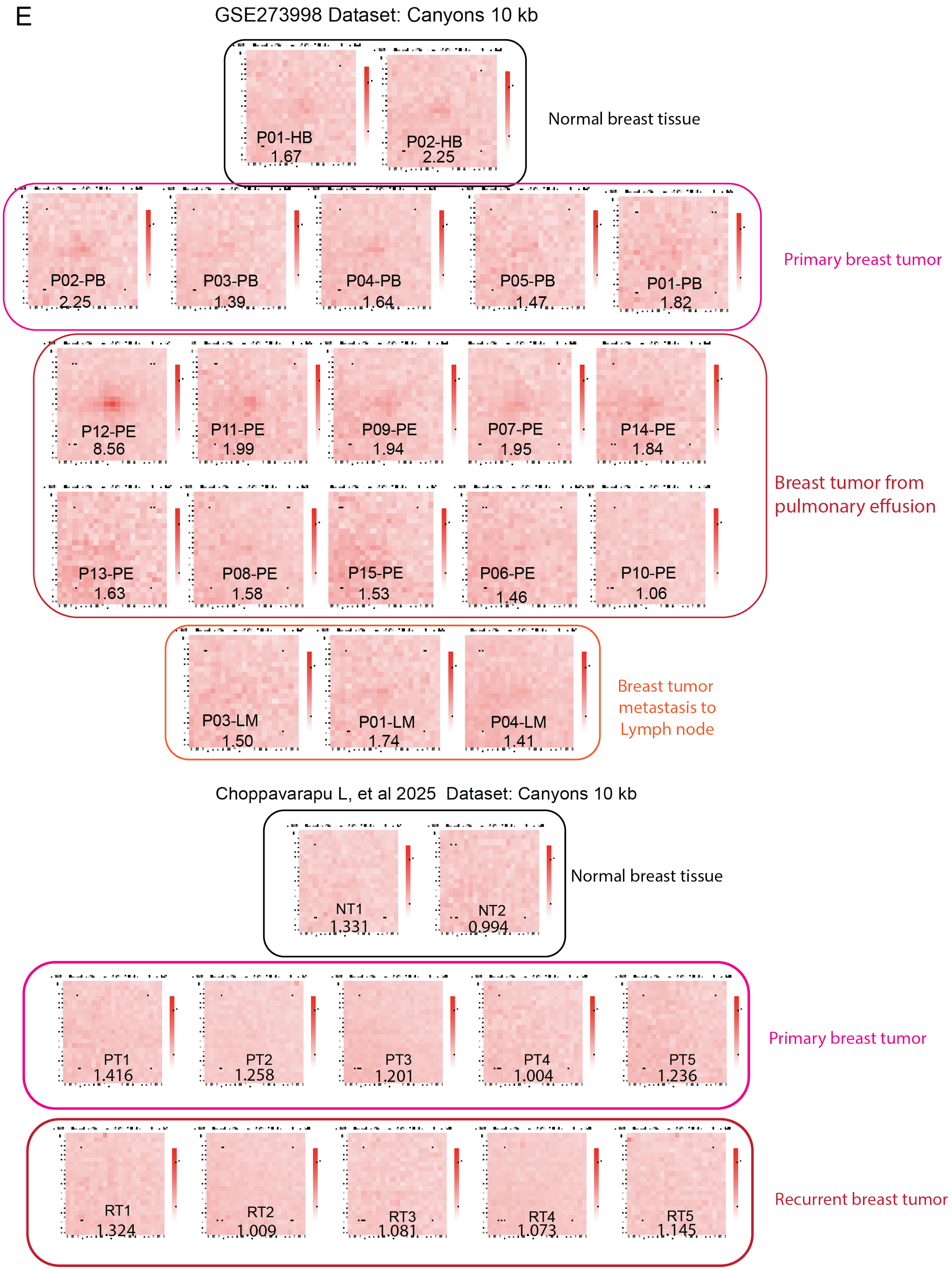
E
GSE273998 Dataset: Canyons 10 kb
P01-HB
1.67
P02-HB
2.25
Normal breast tissue
P02-PB
2.25
P03-PB
1.39
P04-PB
1.64
P05-PB
1.47
P01-PB
1.82
Primary breast tumor
P12-PE
8.56
P11-PE
1.99
P09-PE
1.94
P07-PE
1.95
P14-PE
1.84
P13-PE
1.63
P08-PE
1.58
P15-PE
1.53
P06-PE
1.46
P10-PE
1.06
Breast tumor from pulmonary effusion
P03-LM
1.50
P01-LM
1.74
P04-LM
1.41
Breast tumor metastasis to Lymph node
Choppavarapu L, et al 2025 Dataset: Canyons 10 kb
NT1
1.331
NT2
0.994
Normal breast tissue
PT1
1.416
PT2
1.258
PT3
1.201
PT4
1.004
PT5
1.236
Primary breast tumor
RT1
1.324
RT2
1.009
RT3
1.081
RT4
1.073
RT5
1.145
Recurrent breast tumor

F

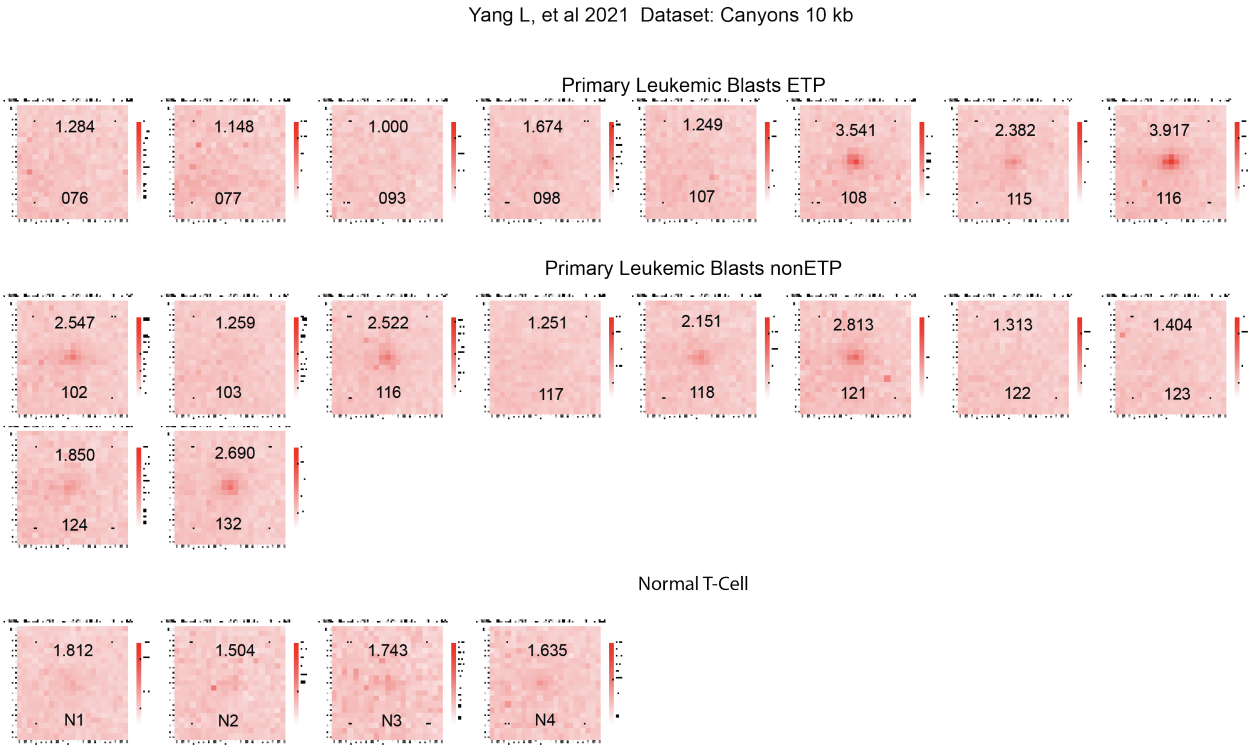


**Supplemental Figure 1. The LPL APA plot of the pan-cancer samples collected in this study.**

**(A).** LPL APA plots and scores of AML samples used in this study. The sample annotation obtained from the dataset is shown. The APA score calculation uses the DNA methylation Grand Canyon regions as the LPL anchor. The samples are ranked by APA score

**(B).** LPL APA plots and scores of pediatric neurological tumor samples used in this study. The sample annotation obtained from the dataset was shown. The APA score calculation uses the DNA methylation Grand Canyon regions as the LPL anchor.

**(C).** LPL APA plots and scores of prostate cancer samples used in this study. The sample annotation obtained from the dataset was shown. The APA score calculation uses DNA methylation Grand Canyon regions as the LPL anchor.

**(D).** LPL APA plots and scores of colon cancer samples used in this study. The sample annotation obtained from the dataset was shown. The APA score calculation uses DNA methylation Grand Canyon regions as the LPL anchor.

**(E).** LPL APA plots and scores of breast cancer samples used in this study. The sample annotation obtained from the dataset was shown. The APA score calculation uses DNA methylation Grand Canyon regions as the LPL anchor.

**(F)**. LPL APA plots and scores of T-ALL samples used in this study.

### Supplemental Figure 2

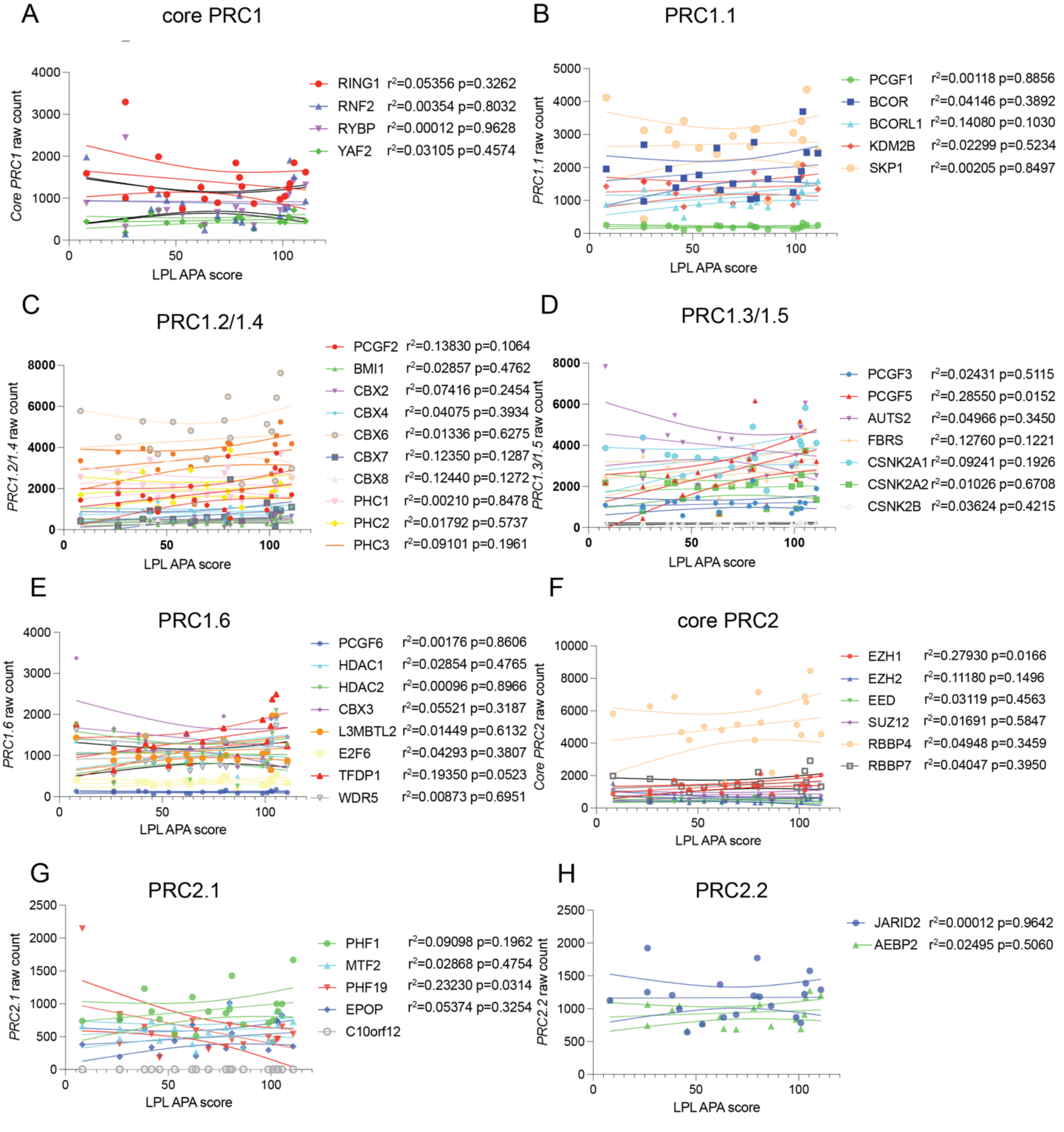


**Supplemental Figure 2. The correlation of Polycomb subunit proteins with LPL APA score in PFA ependymoma.**

**(A).** The Pearson correlation of core PRC1 complex subunit protein's expression with LPL APA score.

**(B).** The Pearson correlation of PRC1.1 complex subunit protein's expression with LPL APA score.

**(C).** The Pearson correlation of PRC1.2/1.4 complex subunit protein's expression with LPL APA score.

**(D).** The Pearson correlation of PRC1.3/1.5 complex subunit protein's expression with LPL APA score.

**(E).** The Pearson correlation of PRC1.6 complex subunit protein's expression with LPL APA score.

**(F).** The Pearson correlation of core PRC2 complex subunit protein's expression with LPL APA score.

**(G).** The Pearson correlation of PRC2.1 complex subunit protein's expression with LPL APA score.

**(H).** The Pearson correlation of core PRC2.2 complex subunit protein's expression with LPL APA score.

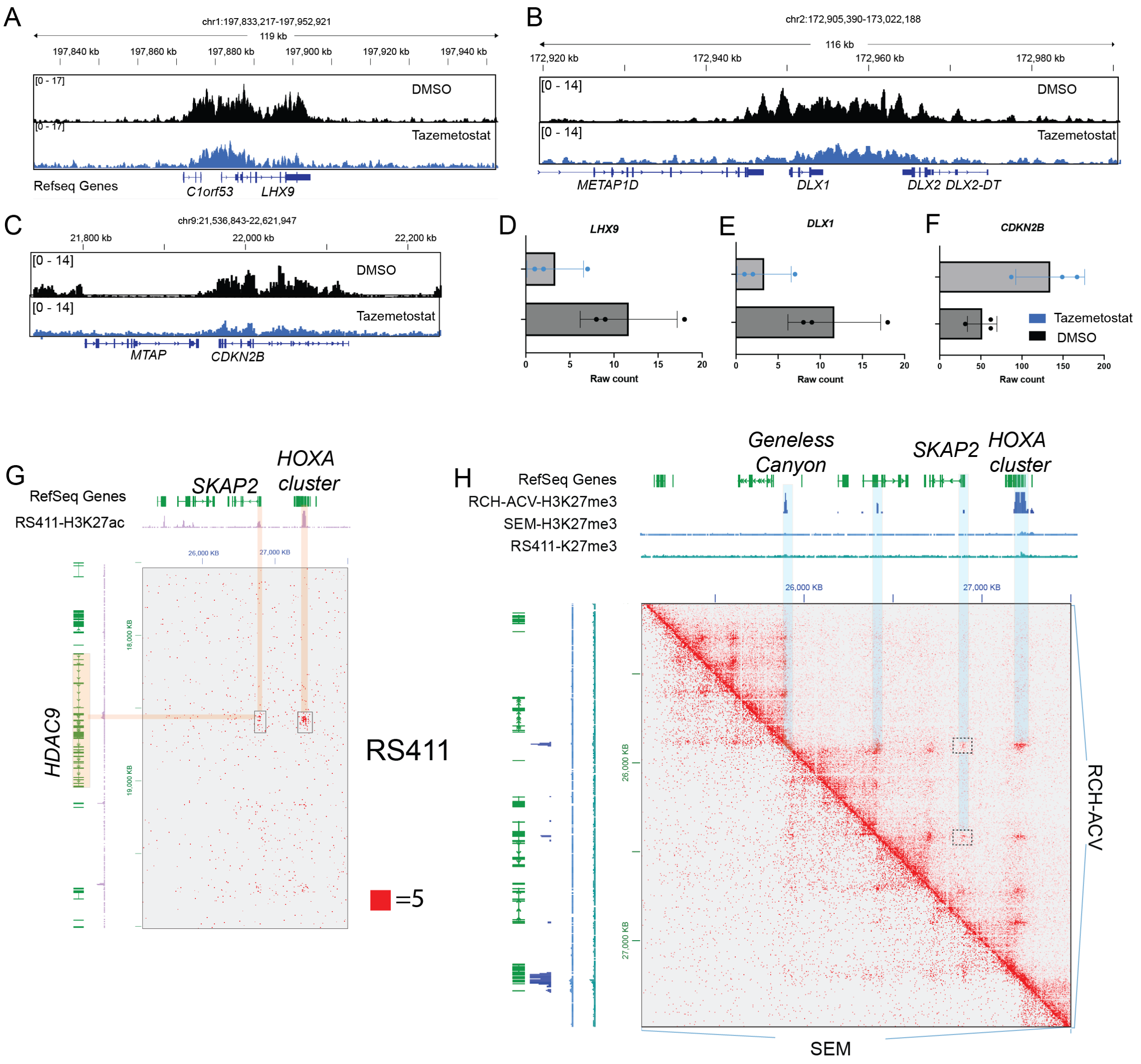


**Supplemental Figure 3. The epigenomic status and gene expression of LPL anchor genes in AML and B-ALL**

**(A-C).** The H3K27me3 distribution on LPL anchor loci in AML5832 cells treated with EZH2 inhibitor and DMSO vehicle control. **(A).** *LHX9* locus **(B).** *DLX1* locus **(C).** *CKDN2B* locus

**(D-F).** The gene expression (raw RNA-seq read counts) of AML5832 cells treated with EZH2 inhibitor and DMSO vehicle control. **(D).** *LHX9* **(E).** *DLX1* **(F).** *CKDN2B*.

**(G).** HiC contact map showing Long range H3K27ac marked interaction between *HOXA* cluster, *SKAP2* and *HDAC9* from 9Mb upstream.

**(H).** HiC contact map showing the ectopically formed H3K27me3 region loops in RCH-ACV. The boxed regions show the interactions between *SKAP2* and *Geneless Canyon.*

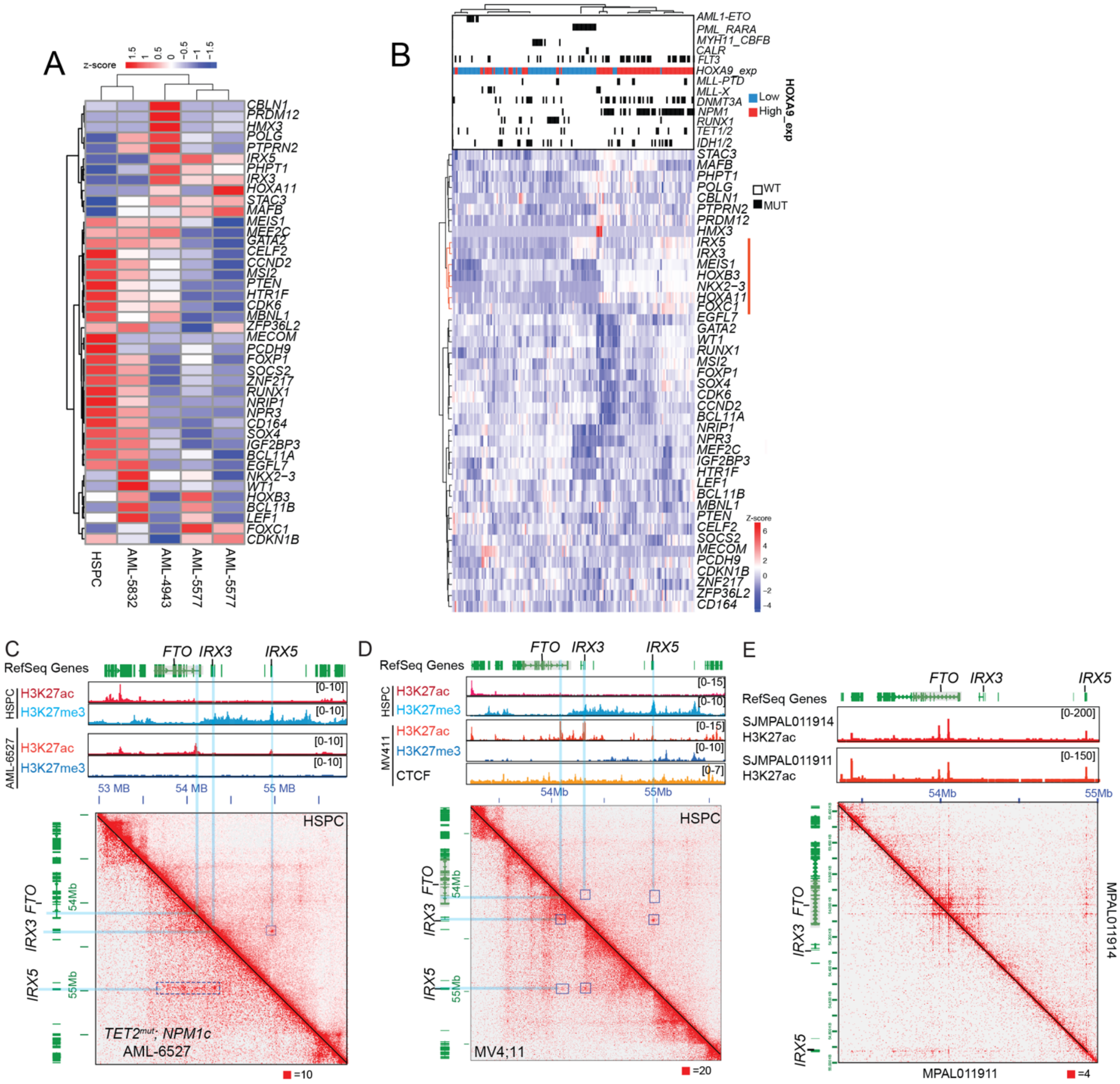

**Supplemental Figure 4. LPL anchor gene is activated in AML and lineage-ambiguous leukemia.**

**(A).** Differentially expressed LPL anchor genes in AMLs used in this study.

**(B).** Differentially expressed LPL anchor genes in TCGA AMLs. A cluster of genes (*IRX3, IRX5*, *NKX2-3*, and *FOXC1*) correlated with *HOXA9* gene expression and NPM1 mutation was labelled.

**(C-D).** HiC contact map of *FTO-IRX3-IRX5* locus in primary AML (**C**) and AML cell lines (**D**). Comparison between HSPC (Upper triangle) and AML (Lower triangle) was shown. *IRX3, IRX5* and *FTO* enhancer were highlighted in blue. Boxed regions show the interaction

**(E).** *FTO-IRX3-IRX5* locus in primary lineage ambiguous leukemia. H3K27ac HiChIP contact map was shown in two samples (MAPL011914 on the upper triangle and MAPL011911 on the lower triangle)
